# Antennal transcriptome analysis and identification of candidate chemosensory genes of the harlequin ladybird beetle, *Harmonia axyridis* (Pallas) (Coleoptera: Coccinellidae)

**DOI:** 10.1101/2021.02.01.429221

**Authors:** Gabriele Rondoni, Alessandro Roman, Camille Meslin, Nicolas Montagné, Eric Conti, Emmanuelle Jacquin-Joly

## Abstract

In predatory ladybirds (Coleoptera: Coccinellidae), antennae are important for chemosensory reception used during food and mate location, and for finding a suitable oviposition habitat. Based on NextSeq 550 Illumina sequencing, we assembled the antennal transcriptome of mated *Harmonia axyridis* (Pallas) (Coleoptera: Coccinellidae) males and females and described the first chemosensory gene repertoire expressed in this species. We annotated candidate chemosensory sequences encoding 26 odorant receptors (including the coreceptor, Orco), 17 gustatory receptors, 27 ionotropic receptors, 31 odorant-binding proteins, 12 chemosensory proteins and 4 sensory neuron membrane proteins. Maximum-likelihood phylogenetic analyses allowed to assign candidate *H. axyridis* chemosensory genes to previously described groups in each of these families. Differential expression analysis between males and females revealed low variability between sexes, possibly reflecting the known absence of relevant sexual dimorphism in the structure of the antennae and in the distribution and abundance of the sensilla. However, we revealed significant differences in expression of three chemosensory genes, namely 2 male-biased odorant-binding proteins and 1 male-biased odorant receptor, suggesting their possible involvement in pheromone detection. Our data pave the way for improving the understanding of the molecular basis of chemosensory reception in Coccinellidae.

**Summary:** The predatory harlequin ladybird *Harmonia axyridis* (Pallas) (Coleoptera: Coccinellidae) has been widely released for classical and augmentative biological control programs of insect herbivores and is now distributed worldwide. Because of its invasive behavior and the threat it can pose to local biodiversity, this ladybird has been adopted as a model species for invasive biocontrol predators. A huge existing literature is available on this species. However, little is known about the mechanisms underlying *H. axyridis* smell and taste, even though these senses are important in this ladybird for courtship, mating and for locating suitable habitats for feeding and oviposition. Here we describe the first chemosensory gene repertoire that is expressed in the antennae of male and female *H. axyridis*. Our findings would likely represent the basis for future functional studies aiming at increasing the efficacy of *H. axyridis* in biological control or at reducing its populations in those areas where the ladybird has become a matter of concern due to its invasiveness.

## 1. Introduction

Chemosensory reception is important in insects, including predacious ladybirds (Coleoptera: Coccinellidae), for food and mate location, and for finding a suitable habitat for oviposition [1–3]. Volatile molecules are typically detected by insects through neurons housed in chemosensory sensilla mainly located on antennae. In these organs, the chemical stimuli are transformed into electrical signals that will be transmitted to the brain [4]. In the proposed process of odorant detection, molecules are first bound and transported by odorant-binding proteins (OBPs) and possibly chemosensory proteins (CSPs) within the sensillum lymph, then detected by odorant receptors (ORs) or/and ionotropic receptors (IRs) expressed at the membrane of olfactory sensory neurons. Other protein families are also involved, such as the sensory neuron membrane proteins (SNMPs) for pheromone detection in some species [4,5]. Apart from olfaction, insect antennae are also involved in taste and numerous antennal transcriptomes have identified gustatory receptors (GRs), among them candidate sugar and CO2 receptors [6]. The function and characteristics of all these gene families in insects have been extensively reviewed, especially in model species such as *Drosophila melanogaster* (Meigen) [4,7–10].

Concerning insects of agricultural economic importance, chemosensory genes have been well described in Lepidoptera and in herbivorous and xylophagous Coleoptera species [e.g., 11–13]. Recent antennal transcriptome analyses focused also on predatory insects, belonging to, e.g., Neuroptera [14,15], Hemiptera [16,17] and Diptera [18]. Surprisingly, despite the worldwide importance of predatory ladybirds (Coleoptera: Coccinellidae) for classical and augmentative biological control [reviewed by 19–21], no detailed information is available at present. Most ladybird species are important natural enemies of various crop pests, such as aphids, scales, whiteflies or mites [22,23]. In particular, *Harmonia axyridis* (Pallas) is the most studied ladybird. This species is native from Asia and, starting from 1916 in the USA, it was released worldwide for biological control of herbivore pests. Currently, it is present in more than 38 countries [reviewed by 24]. Besides the beneficial role that *H. axyridis* exhibits in pest suppression [25], this ladybird raises concerns on the possible negative effects it may cause to the invaded community of predators, through intraguild predation [e.g., 26–28] and indirect competition [29,30].

Behavioural and electrophysiological experiments demonstrated that *H. axyridis* responds to volatile semiochemicals [1,31,32]. Additionally, various types of antennal chemoreceptor sensilla (notably basiconica, chaetica and grooved peg) have been recently characterized [33]. Therefore, understanding the molecular basis of *H. axyridis* chemosensory reception, in particular olfaction, is likely to provide new information to increase the efficacy of this predator in biological control or to reduce its populations in those areas where *H. axyridis* has become a concern for local biodiversity [34]. In this study, we conducted a transcriptomic analysis of *H. axyridis* adult antennae, identified candidate chemosensory genes and investigated their differential expression between males and females. Additionally, we constructed phylogenetic trees and inferred the evolutionary relationships of putative *H. axyridis* chemosensory genes with other coleopteran species.

## 2. Materials and Methods

### 2.1 Insect rearing and antenna collection

A culture of *H. axyridis* was settled in the laboratory from adults collected in Central Italy (Perugia Province). Adults and larvae were reared in plastic cylindrical cages (Ø = 25 cm, height = 30 cm) covered by a fine tissue mesh to allow ventilation. Broad bean plants moderately infested with *Aphis fabae* Scop. were used as a food source for larvae and adults. Plants were changed every 2–3 days and the egg batches were collected and isolated in a new cage for hatching. For tissue collection, and to control adult age and mating status, newly emerged adults were isolated for 3 days (aphids provided) for completing their pigmentation and maturation (pheromone production), then sexed. Groups of 20 males and 20 females were paired in small net cages (30 cm × 30 cm × 30 cm). Each cage contained an aphid-infested plant, to allow mating. Plants were replaced every two days. Mating was observed and confirmed by the presence of physogastric females. After 4 to 6 days, insects were collected for dissection of the antennae under a stereomicroscope. Antennae were immediately stored in a 2mL Eppendorf tube constantly immersed in liquid nitrogen and stored at −80°C until RNA extraction. For each replication, collection of the antennae was conducted daily from 10:00 to 15:00 and split between different days. For each sex, three biological replications were conducted in total, each consisting of 200 antennae collected from ~100 individuals.

### 2.2 RNA purification, cDNA library preparation and sequencing

Total RNA was obtained using RNeasy Plus Mini Kit (Qiagen, USA) following the manufacturer’s protocol for animal samples [35]. The RNA concentration was determined using Qubit Fluorometer and Qubit RNA BR Assay (Life Technologies). RNA integrity was determined using Fragment Analyzer (Agilent Technologies). Libraries were prepared using Illumina TruSeq Stranded mRNA according to the sample preparation guide (Part #15031047, Rev. E, October 2013) for Illumina paired-end indexed sequencing. The resulting libraries were validated using Fragment Analyzer to check size distribution. Concentration of library samples was defined based on Qubit Fluorometer quantification and average library size. Indexed DNA libraries were normalized to 4 nM and then pooled in equal volumes. The pool was loaded at a concentration of 1.1 pM onto an Illumina NextSeq 550 Mid Output Flowcell (with 1% of PhiX control). The samples were then sequenced using the Illumina V2, 2×75 bp paired-end run. Library preparation and sequencing were conducted by Polo Genetica, Genomica e Biologia (Polo GGB, Siena, Italy). The raw data from Illumina sequencing were deposited in the NCBI Short Read Archive (SRA) database (BioProject ID PRJNA698239).

### 2.3 Assembly and functional annotation

De-novo assembly was performed within Galaxy [36] framework. An initial assessment of the quality of the raw reads in fastq format was conducted using FastQC [37]. Reads were then converted in fastqsanger format with FastQ Groomer [38] and low-quality reads were trimmed using Trimmomatic v. 0.32.3 [39] with a required average quality value of 20 and a minimum length of reads to be kept of 30. Identification and removal of rRNA-like sequences were conducted using riboPicker v. 0.4.3 [40]. De-novo assembly was conducted with Trinity [41] by setting ‘Reverse-Forward’ strand-specific library type, a minimum contig length of 200 and a minimum count for K-mers to be assembled of 1. TransDecoder [41] was used to identify coding regions. Redundant sequences were removed using CD-HIT-EST [42] with a 98% sequence identity threshold, 8 word-length size and default options. Predicted peptides were annotated using the BLASTp algorithm (1e^−5^ threshold) on the Swiss-Prot protein database. Mapping, annotation and InterPro analysis were conducted using Blast2GO within OmicsBox v1.4.11. Molecular function, biological process and cellular component were derived for each gene [43]. The quality of the assembled transcriptome (e.g., total number of contigs, total contigs ≥ 500 bp, largest contig, N50) was evaluated with QUAST [44]. BUSCO was used to verify the completeness of the assemblage [45].

### 2.4 Identification of differentially expressed transcripts

Bowtie [46] alignment method and RSEM were used to align the reads on the transcriptome and calculate the raw read numbers and TPM (transcripts per kilobase million, table S3) expression value [47]. Results were analyzed using edgeR within *edgeR* package in R [48], adopting a significance level of false discovery rate (FDR) = 0.10 [as in 49,50]. Transcripts that exhibited a low expression (0.25 counts per million reads) were filtered out from the analysis [similar to 51]. The number of total reads was normalized according to the trimmed mean of M values (TMM) [52].

### 2.5 Identification of chemosensory genes and phylogenetic analyses

Chemosensory genes (~4,000 sequences) collected from NCBI and published literature were used as queries in tBLASTn (1e^−5^ E-value) against cleaned trinity assemblage. For each transcript, the best hit was considered. Protein domains were predicted using TMHMM 2.0 [53] and SignalP 3.0 [54]. The presence and number of conserved cysteine residues in candidate OBPs and CSPs were visually assessed [55]. Phylogenetic trees were constructed using *H. axyridis* candidate chemosensory proteins and proteins from other Coleoptera families, including those closely related to ladybirds [56]. Considered species were *Tribolium castaneum* (Herbst) (Tenebrionidae) [57–59], *Anoplophora glabripennis* (Motschulsky) (Cerambycidae) [60,61], *Dendroctonus ponderosae* (Hopkins) (Curculionidae) [62]*, Ambrostoma quadriimpressum* (Motschulsky), *Onthophagus taurus* (Schreber) and *Anomala corpulenta* (Motschulsky) (Scarabaeidae), and *Phyllotreta striolata* (F.) (Chrysomelidae) [63–65]. Functionally characterized odorant receptors from the beetles *Megacyllene caryae* (Gahan.), *Ips typographus* L. and *Rhyncophorus ferrugineus* (Olivier) were included [66–69] as well as gustatory receptors from *D. melanogaster* [13]. Similarly, sequences of functionally characterized odorant- and pheromone-binding proteins were considered from beetles, i.e., *Anomala cuprea* (Hope) and *Anomala octiescostata* Burmeister [70], *Agrilus mali* Matsumura [71], *Cyrtotrachelus buqueti* Guérin-Méneville [72], *Holotrichia oblita* Faldermannor [73,74], *Holotrichia parallela* (Motschulsky) [75], *Phyllopertha diversa* Waterhouse [76], *Popillia japonica* Newman [77], *R. ferrugineus* [78], and from the aphid *Acyrthosiphon pisum* (Harris) [79]. Sequences were aligned using MAFFT v.7 with Auto (for ORs, GRs and IRs) or FFT-NS-i (for OBPs, CSPs and SNMPs) iterative refinement methods, and default parameters [80]. Maximum Likelihood phylogenies were built using PhyML 3.0 [81], using the substitution model JTT and a SH-like approximate likelihood-ratio test for node support estimation [9,82]. Trees were viewed and edited using FigTree v1.4.0 (http://tree.bio.ed.ac.uk/software/figtree/) and CorelDRAW X3. A phylogenetic analysis was conducted with candidate *H. axyridis* ORs and GRs and known ORs and GRs from *T. castaneum*, to verify effective phylogenetic separation between members of the two chemosensory receptor families [83]. In two cases, sequences that were originally named as ORs were later assigned to GRs (HaxyGR16 and HaxyGR17). Concerning the nomenclature, the OR co-receptor was named as “Orco”. Sequences that clustered with non-chemosensory ionotropic glutamate receptors of *D. melanogaster* [84] were excluded from the final phylogenetic tree analysis and from the list of HaxyIRs. When multiple HaxyIR sequences clustered together, they were aligned to verify that they are not part of the same transcript. Additionally, we referred to *D. melanogaster* and *T. castaneum* for HaxyIRs that clustered with known IRs [table S3 in 84]. Six candidate chemosensory proteins that exhibited high identity with ejaculatory bulb proteins and presented six conserved cysteine (C) residues were excluded. SNMPs were classified according to SNMP1 and SNMP2 groups.

## 3. Results

### 3.1 Transcriptome sequencing, assembly, and identification of chemosensory genes

Sequencing returned approximatively 55 million raw reads. The accuracy of Q30 base call was 92.6% (table S1.1). The total assembly (males + females) returned 66,728 contigs, with the largest contig of 16,954 bp and a N50 of 1,971 bp (table S1.2). The total number of contigs ≥ 500 bp in length was 30,600. Gene coverage was high, as the assembled transcriptome contained 93.2% complete BUSCO groups (single-copy: 66.3%; duplicated: 26.9%), 2.2% fragmented and 4.6% missing BUSCO genes. Searches against the Swiss-Prot database returned 27,698 transcripts showing sequence similarity to known proteins (reported in table S2). Additionally, 28,991 and 35,318 transcripts were assigned with 1 or more GO terms or InterPro IDs, respectively. Most abundant biological processes and molecular functions were related to cellular and metabolic functions as well as chemosensory processes (“response to stimulus”, “localization” and “protein binding”) (figure S1). Bioinformatics analyses identified a total of 117 unigenes from *H. axyridis* transcriptome that belonged to gene families putatively involved in insect chemoreception: 26 ORs, 17 GRs, 27 IRs, 31 OBPs, 12 CSPs, 4 SNMPs (tables S5 to S11). Expression analysis revealed 133 contigs differentially expressed between males and females (tables S3 and S4), including three chemosensory gene isoforms. Notably, two candidate OBPs (HaxyOBP11 and HaxyOBP27, table S8) were highly expressed in males compared to females (FDR < 0.05). In details, HaxyOBP11 exhibited 82.6% amino acid identity with OBP2 of *Cryptolaemus montrouzieri*, and HaxyOBP27 87.6% with OBP38 of *Holotrichia parallela*. One OR (HaxyOR5, Table S5) was more expressed in males compared to females (FDR = 0.086) and exhibited 57% amino acid identity with OR49b-like of *Leptinotarsa decemlineata*.

### 3.2 Odorant receptors (ORs)

Of the 26 HaxyORs, full-length ORFs were identified for 11 of them, with lengths ranging from 339 to 480 amino acids and 4 to 7 predicted transmembrane domains (table S5). The remaining 15 ORs corresponded to partial sequences, encoding 102 to 410 amino acids. Except for Orco, which is highly conserved between insect families, candidate HaxyORs exhibited low amino acid identity with ORs from other coleopteran species. Phylogenetic analysis revealed that the different ORs are included in 4 out of 9 groups known for Coleoptera (figure 1) [9]. In details, seven candidate HaxyORs clustered in group 2A, three HaxyORs clustered in group 3, six HaxyORs clustered in group 5A, and nine HaxyORs, including the male-biased HaxyOR5 (see section 3.1) belonged to group 7. No candidate OR was detected in groups 1, 2B, 4, 5B and 6. HaxyORs that grouped in the same clades 2A, 3, 5A or 7 were aligned to check whether these fragments correspond to different fragments of the same protein or are different proteins. Alignments did not reveal conserved overlapping regions thus suggesting they are different ORs.

**Figure 1.**
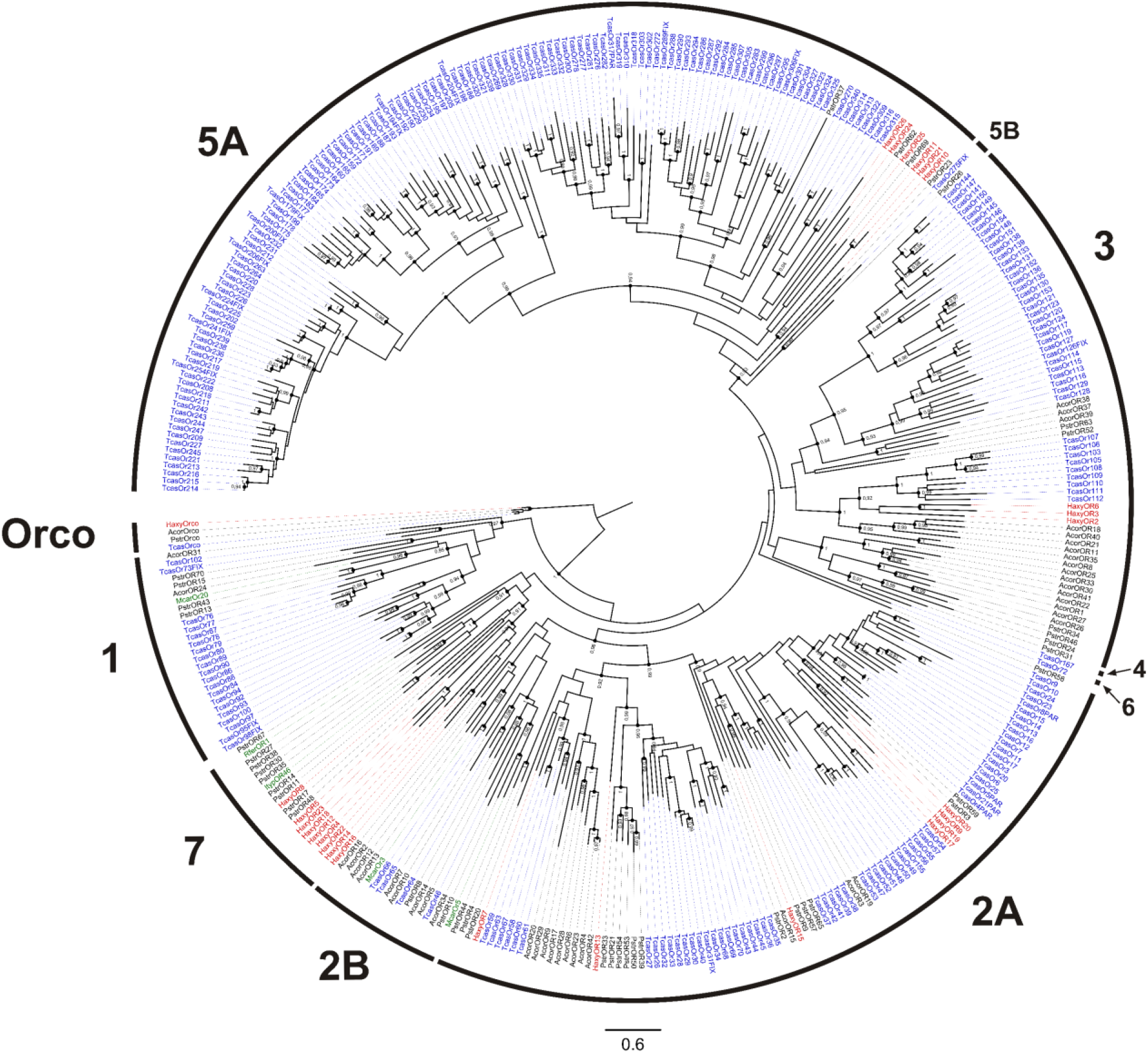
Phylogenetic tree of ORs. Red: *Harmonia axyridis* (Haxy); blue: *Tribolium castaneum* (Tcas); black: *Anomala corpulenta* (Acor) and *Phyllotreta striolata* (Pstr); green: functionally characterized ORs of *Megacyllene caryae* (Mcar), *Ips typographus* (Ityp), and *Rhyncophorus ferrugineus* (Rfer). External numbers represent known groups of ORs. Numbers and symbols at nodes represent support values higher than 0.9, where 1 represents maximal support.

### 3.3 Gustatory receptors (GRs)

Seventeen contigs encoding candidate GRs were identified in the transcriptome (table S6). Of them, only two encoded full-length proteins (392 and 440 amino acids). HaxyGR11, HaxyGR12, and HaxyGR13 grouped with *D. melanogaster* GR21a and HaxyGR15 grouped with *D. melanogaster* GR63a (figure 2). In *D. melanogaster*, these two genes are responsible for CO2 detection [6]. HaxyGR1, HaxyGR3, HaxyGR4 and HaxyGR8 grouped with *Drosophila* GRs known to be involved in sugar detection [85]. As for ORs, we aligned amino acid sequences of the different fragmented GRs to ensure they represent different proteins. Inspection of the alignment for HaxyGR1, HaxyGR4 and HaxyGR8 suggested that these fragments are indeed different GRs. Conversely, alignment of HaxyGR12, HaxyGR11 and HaxyGR13 revealed overlapping regions (5-7 amino acids), thus suggesting that the three fragments may be part of the same protein. HaxyGR2 and HaxyGR6 clustered with TcasGR20 from *T. castaneum*, which is a receptor for mannitol and sorbitol [86]. No GR appeared to be differentially expressed between sexes.

**Figure 2.**
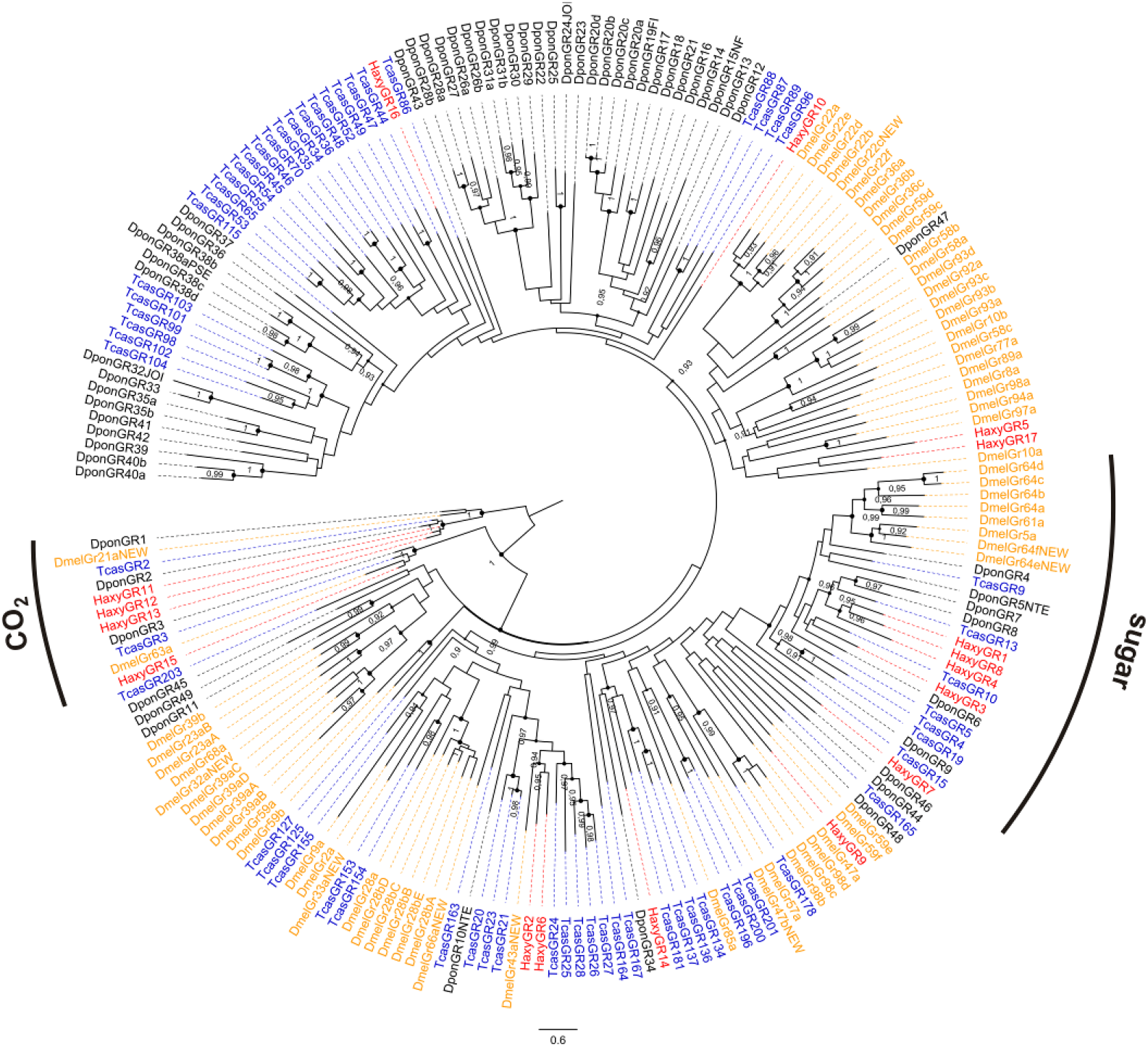
Phylogenetic tree of GRs. Red: *Harmonia axyridis* (Haxy); blue: *Tribolium castaneum* (Tcas); orange: *Drosophila melanogaster* (Dmel); black: *Dendroctonus ponderosae* (Dpon). Numbers and symbols at nodes represent support values higher than 0.9, where 1 represents maximal support.

### 3.4 Ionotropic receptors (IRs)

Twenty-seven IRs were identified in total, 5 of them encoding full-length proteins, from 301 to 938 amino acids (table S7). Phylogenetic tree (figure 3) revealed the presence of HaxyIR orthologues of *D. melanogaster* and *T. castaneum* IRs, such as the IR coreceptors IR8a and IR25a [87]. Alignments did not reveal conserved overlapping regions between sequences. The high number of IRs found in *H. axyridis* can be explained by the conservation of IRs across insect families, making them easy to identify by sequence homology, contrary to ORs and GRs that are usually more divergent. Additionally, we found impressive IR duplications in the coreceptor IR25a (7 sequences) and IR8a (3 sequences) subfamilies. All are non-overlapping fragments, but we cannot exclude that some could correspond to the same IR, considering that IR genes are usually very large. No IR appeared to be differentially expressed between sexes.

**Figure 3.**
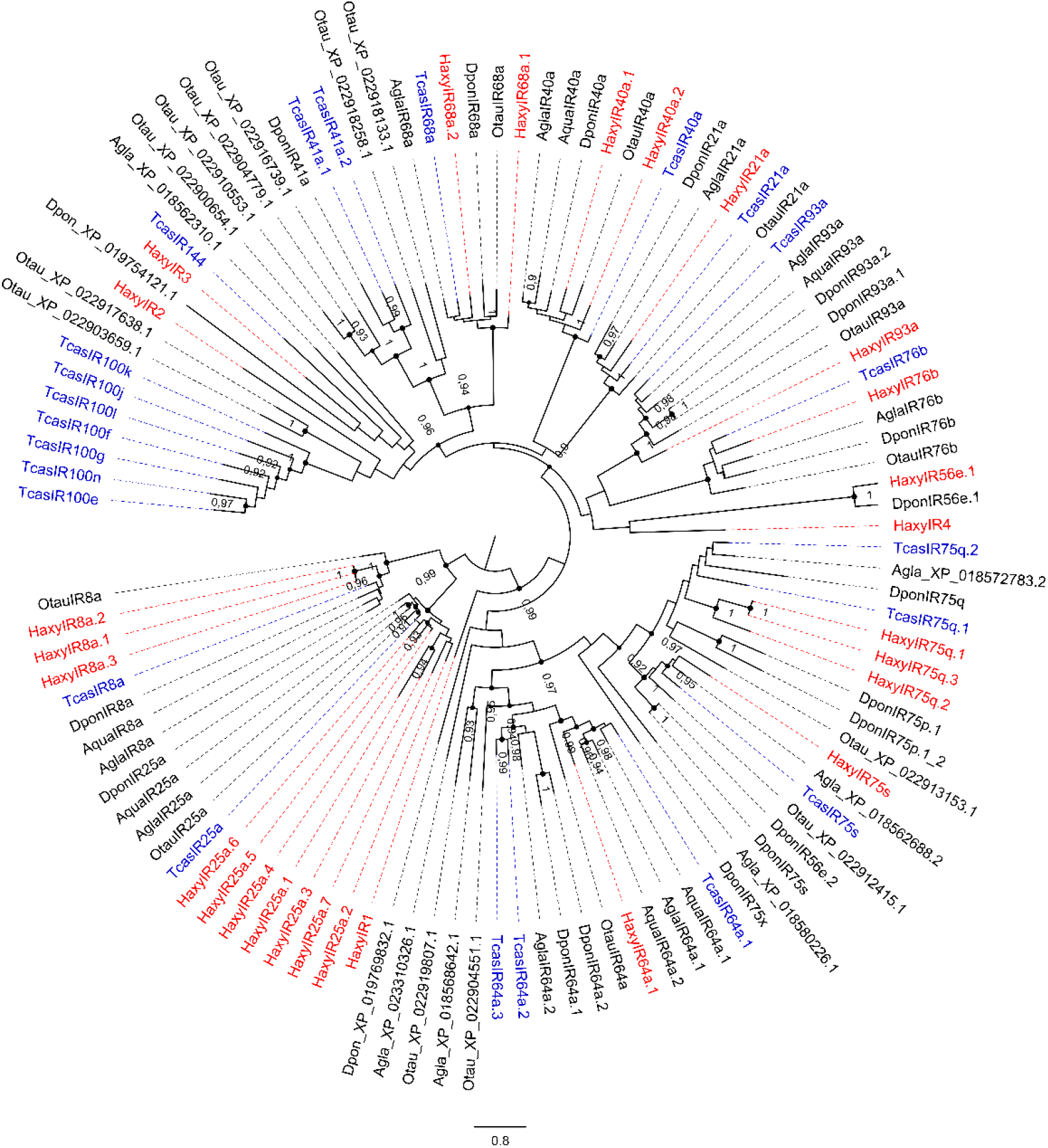
Phylogenetic tree of IRs. Red: *Harmonia axyridis* (Haxy); blue: *Tribolium castaneum* (Tcas); black: *Ambrostoma quadriimpressum* (Aqua), *Anoplophora glabripennis* (Agla), *Dendroctonus ponderosae* (Dpon), and *Onthophagus taurus* (Otau). Numbers and symbols at nodes represent support values higher than 0.9, where 1 represents maximal support.

### 3.5 Odorant-binding proteins (OBPs)

In total, 31 OBPs were identified in the transcriptome of *H. axyridis*. For 16 of them, the complete ORF was detected, with a length varying from 123 to 265 amino acids (table S8). Twenty-four HaxyOBPs presented a predicted signal peptide. Twenty of the candidate OBPs belonged to the classic-OBP family, exhibiting the 6 conserved cysteine (C) residues representative of this OBP subfamily. Ten OBPs belonged to the minus-C OBP subgroup with 4 conserved cysteine residues. One OBP (HaxyOBP26) belonged to the plus-C subfamily and clustered with OBP5E of *T. castaneum* (figure 4). The complete HaxyOBP30 clustered with TcasOBP8B, an OBP that exhibits a non-conserved cysteine pattern, with extra amino acids between C1 and C2 [57]. Similarly, HaxyOBP30 exhibited non-conventional interval between C1 and C2. Both HaxyOBP11 and HaxyOBP27 were more expressed in males. HaxyOBP11 clustered with 3 other OBPs from *H. axyridis* and with 4 OBPs from *T. castaneum*. Conversely, it was not possible to find orthologues for HaxyOBP27. Interestingly, HaxyOBP10 and HaxyOBP20 clustered with HoblOBP3 and HoblOBP4, two proteins that bound plant-related compounds, and AmalOBP3, a protein that showed binding affinity with alcohols, esters, terpenoids and 12-15 carbon aldehydes [71,88].

**Figure 4.**
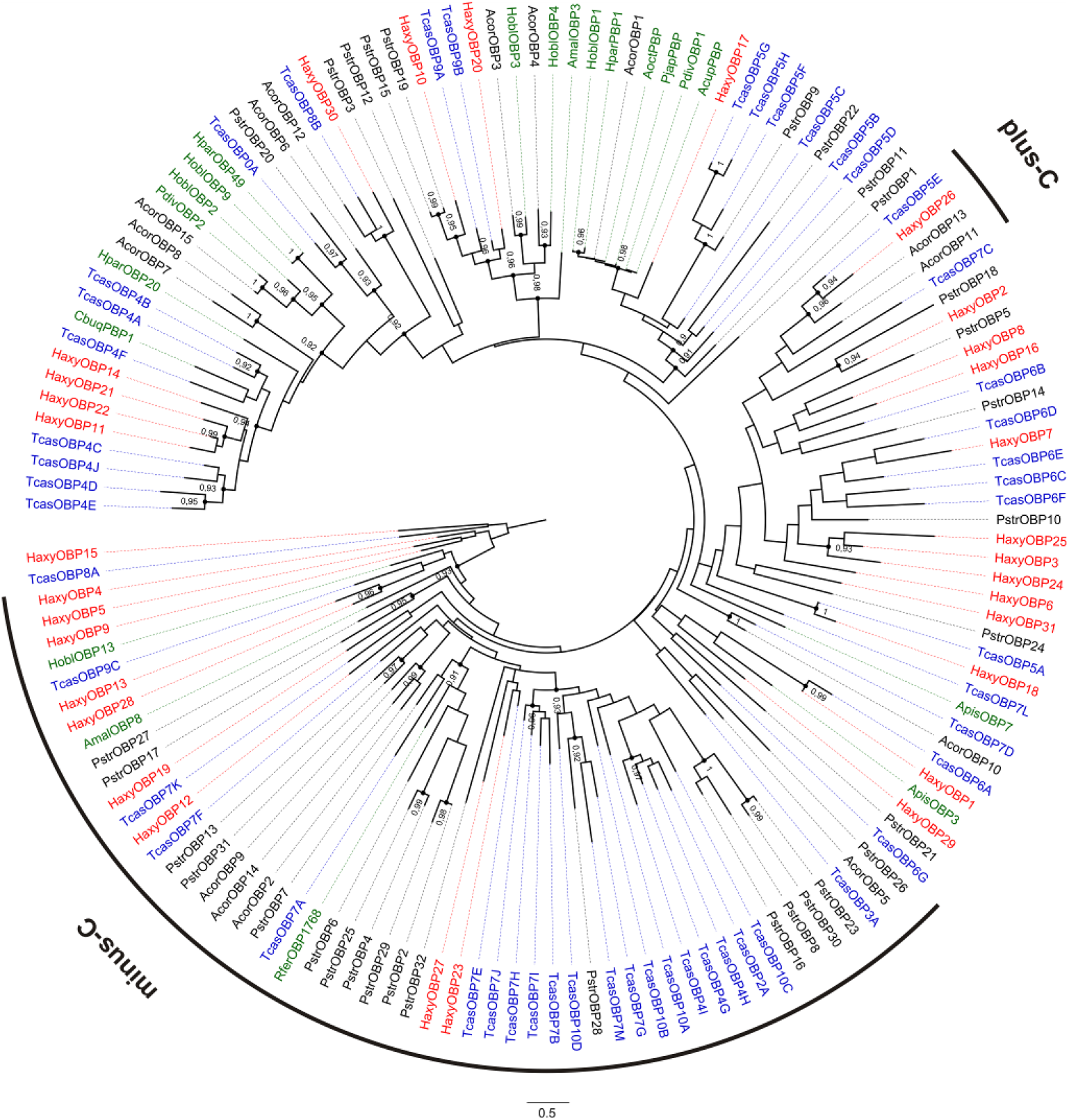
Phylogenetic tree of OBPs. Red: *Harmonia axyridis* (Haxy); blue: *Tribolium castaneum* (Tcas); black: *Anomala corpulenta* (Acor) and *Phyllotreta striolata* (Pstr); green: functionally characterized OBPs and pheromone binding proteins (PBPs) from *Acyrthosiphon pisum* (Apis), *Anomala cuprea* (Acup), *Agrilus mali* (Amal), *Anomala octiescostata* (Aoct), *Cyrtotrachelus buqueti* (Cbuq), *Holotrichia oblita* (Hobl), *Holotrichia parallela* (Hpar), *Phyllopertha diversa* (Pdiv), *Popillia japonica* (Pjap), *Rhynchophorus ferrugineus* (Rfer). Numbers and symbols at nodes represent support values higher than 0.9, where 1 represents maximal support.

### 3.6 Chemosensory proteins (CSPs)

Of the 12 contigs encoding candidate CSPs, 8 encoded full-length proteins (with length from 101 to 153 amino acids). Nine HaxyCSPs possessed a predicted signal peptide and all presented the highly conserved four-cysteine profile (table S9, figure 5). No CSP appeared to be differentially expressed between sexes.

**Figure 5.**
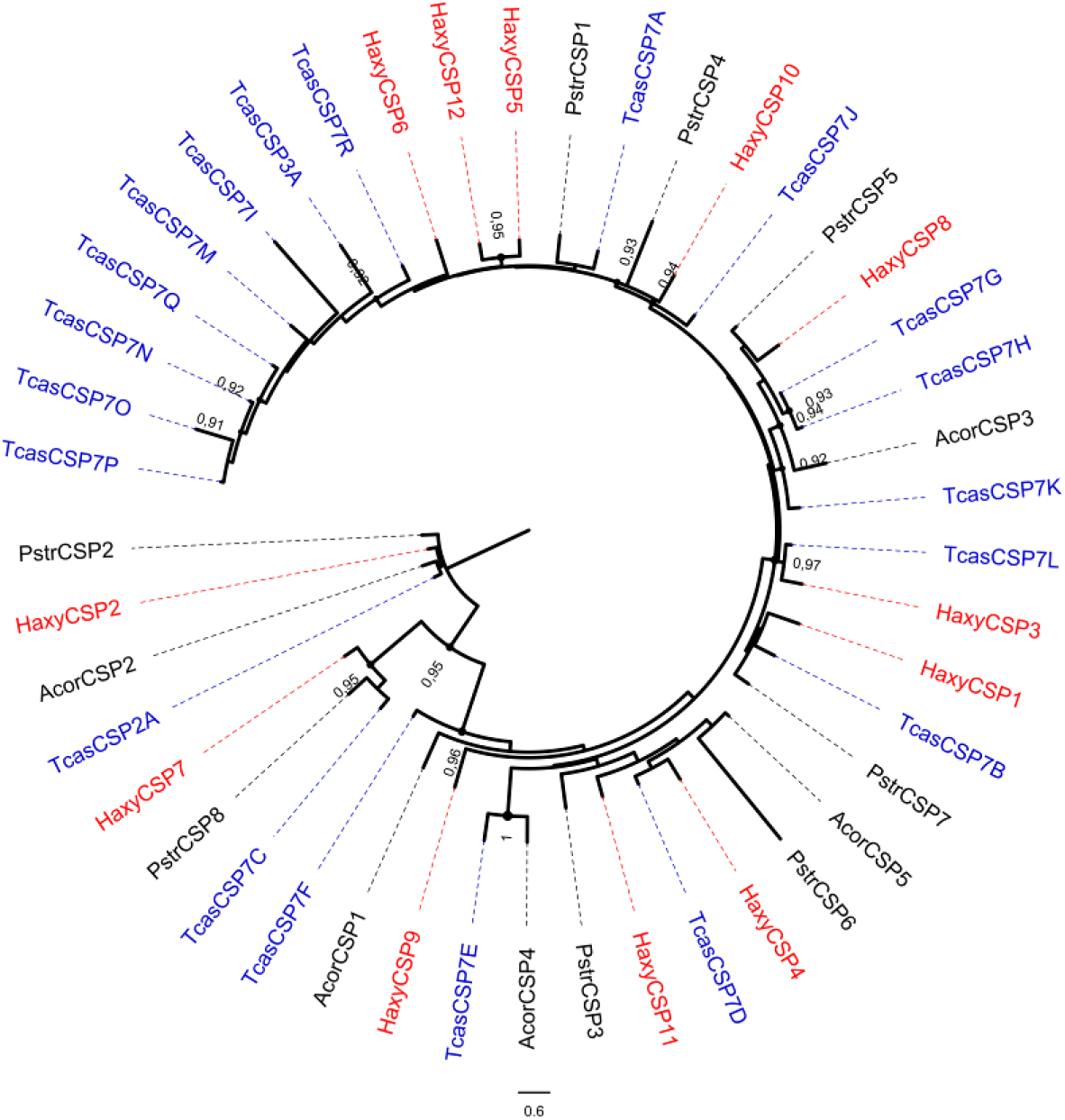
Phylogenetic tree of CSPs. Red: *Harmonia axyridis* (Haxy); blue: *Tribolium castaneum* (Tcas); black: *Anomala corpulenta* (Acor) and *Phyllotreta striolata* (Pstr). Numbers and symbols at nodes represent support values higher than 0.9, where 1 represents maximal support.

### 3.7 Sensory neuron membrane proteins (SNMPs)

Four sequences encoding incomplete SNMPs were identified in the *H. axyridis* transcriptome and belonged to the SNMP1 and SNMP2 insect lineages (table S10, figure S2). HaxySNMP1.1 exhibited 62.5% identity with *T. castaneum* SNMP1, while HaxySNMP2.1 was 63.6% identical to *T. castaneum* SNMP2. No SNMP appeared to be differentially expressed between sexes.

## 4. Discussion

Despite the huge amount of ladybird species that have been released in 130 years of biological control activity, the chemical ecology of this group and the molecular basis of its olfaction are still largely unknown. Here, we identified 117 chemosensory genes expressed in *H. axyridis* adult antennae and provided a first survey of their differential expression between males and females. Most chemosensory genes did not exhibit sex-specific nor sex-biased expression. This is consistent with recent description of antennal morphology that did not reveal relevant sexual dimorphism in *H. axyridis*, neither for the antennal general structure, nor for the types of sensilla and their abundance [33]. However, this does not preclude that the significance of the perceived odors may differ between males or females. Notably, it is known from previous studies that *H. axyridis* males and females respond differently to odors in wind tunnel [89] or electroantennography [90].

The number of ORs we described in *H. axyridis* was a bit lower compared to what has been identified in other beetle transcriptomes, e.g., *I. typographus* (43) and *D. ponderosae* (49) [62,91]. This difference may be related to the different chemical ecologies of these species, as previously suggested [8]. Alternatively, it is possible that we missed some HaxyORs, as our transcriptome was sequenced from adult antennae only (no larvae). However, antennal transcriptomes from other predatory insects exhibited variable numbers of ORs. For instance, only 15 ORs could be identified in the antennal transcriptome of *Cyrtorhinus lividipennis* Reuter (Hemiptera: Miridae) [17], 14 ORs in *Chrysopa pallens* (Rambur) [14] and 37 ORs in *Chrysoperla sinica* (Neuroptera: Chrysopidae) [15], 51 ORs in *Episyrphus balteatus* DeGeer and 42 ORs in *Eupeodes corollae* Fabricius (Diptera: Syrphidae) [18], and 38 ORs in *Arma chinensis* (Fallou) (Hemiptera: Pentatomidae) [92]. Hopefully, further identification of OR genes in other predatory insect antennae will permit to increase coverage of probable unknown putative OR genes in these insects. Additionally, annotation of *H. axyridis* ORs in this species genome will likely reveal the presence of more ORs [93], as it happened for other species [94], although their effective expression will remain to be investigated. Pheromonal blend has been recently characterized for *H. axyridis*, with (–)-β-caryophyllene being the most abundant molecule [95]. The pheromone is emitted by virgin and mated females, primarily under the presence of adequate aphid preys [96]. Major sex pheromone compounds are individually perceived by both sexes, while the complete pheromonal blend elicited behavioral attraction on males only [90]. Possibly, high expression of one OR, HaxyOR5, in males may suggest a role in the detection of one sex pheromone molecule. In support of this latter hypothesis is that HaxyOR5 clustered in the clade 7 of the Coleoptera OR phylogeny, in which pheromone receptors from other Cucujiformia (*R. ferrugineus* and *I. typographus*) have been characterized [68,69]. Together with the fact that Coleoptera ORs exhibit high sequence divergence, this makes it difficult to predict HaxyOR functions. That is why it would be necessary to identify ligands experimentally in the future [reviewed by 7]. Our transcriptomic analysis revealed that 4 candidate sugar receptors were expressed in *H. axyridis* antennae. Sugar is an important component in the diet of predatory ladybirds, notably *H. axyridis* [97]. For instance, when aphid preys are unavailable, *H. axyridis* maintains its presence in the agroecosystem by feeding upon extrafloral nectars [98]. Sugar is also present in aphid honeydew, which may represent an alternative food source for *H. axyridis* [99]. Additionally, HaxyGR2 and HaxyGR6 clustered in the phylogeny with TcasGR20, a receptor which is responsible in *T. castaneum* for mannitol and sorbitol detection [86]. Although it is thought that ladybird maxillary palps are the most adequate structures for the perception of nonvolatile molecules [100], our results suggest a possible role of antennal contact in sugar detection. Possibly, the long sensilla chaetica that are positioned in *H. axyridis* at the tip of the antenna may be involved in the perception of such nonvolatile molecules [33]. Our analysis also revealed the presence of candidate receptors for CO_2_ in *H. axyridis* antennae. Sensilla basiconica are responsible for carbon dioxide detection in *D. melanogaster* [12]. These sensilla are present and abundant in male and female *H. axyridis* antennae [33]. Although this kind of sensilla is known to be involved in the detection of a plethora of molecules in insects, including volatile molecules, it is possible that CO_2_ detection is mediated by such sensilla in *H. axyridis*. The direct effect of artificial changes of CO_2_ concentration on the behaviour of biocontrol agents is still under investigation [101]. To date, it has been shown that CO_2_ has weak or null influence on *H. axyridis* development and feeding behaviour [102]. Concerning IRs, the fact that more than one gene have been found clustering in the IR25a clade was unexpected. A possible explanation is that these fragments are indeed part of one large IR gene that is particularly difficult to reconstruct with our sequencing approach [88]. Alternatively, they may result from IR25a duplication in *H. axyridis*. Gene duplication or pseudogenization of IR25a is rare, but has been observed in some Hymenopteran species [103 and references within], and might be relevant for the acquisition of new functions.

Odorant-binding proteins are important carriers of chemosensory cues, including alarm, aggregation, and sex pheromones. In *A. pisum*, ApisOBP3 and ApisOBP7 are known to carry the alarm pheromone, (E)-β-Farnesene in adults and nymphs [79]. HaxyOBP1 clustered with ApisOBP3, although the aLRT-SH branch support value (0.87) was a bit lower than the adopted threshold of 0.9. *Harmonia axyridis* has been previously shown to respond to (E)-β-Farnesene [90], hence, we can hypothesize for HaxyOBP1 a possible role in binding the aphid alarm pheromone. The male-biased expression of HaxyOBP11 and HaxyOBP27 we observed in this study, and taken into account that sex pheromone is released by *H. axyridis* females, led us to hypothesize that HaxyOBP11 and HaxyOBP27 would play a role as carriers of the female-produced sex-pheromone. In addition, HaxyOBP27 belonged to the minus-C OBP subfamily, as the pheromone-binding protein RferOBP1768 from *R. ferrugineus* does [78]. HaxyOBP18 and HaxyOBP30 clustered with TcasOBP5A and TcasOBP8B respectively. These two *T. castaneum* OBPs have been previously shown to cluster with two *D. melanogaster* OBPs (DmelOBP59a and DmelOBP73a), which are very conserved across insect orders [88]. Hence, it is possible to hypothesize common functions of the two proteins for different insect groups. Finally, the fact that HaxyOBP10 and HaxyOBP20 clustered with *A. mali* and *H. oblita* OBPs that have been shown to have binding affinity for plant volatiles led us to hypothesize a possible involvement of the two proteins in the detection of adequate habitat cues. Our analysis detected four SNMP sequences that belonged to SNMP1 and SNMP2 subfamilies. Although most insects have two SNMPs, there are beetle species with antennal expression of 3 (*I. typographus* and *D. ponderosae*), 4 (*Rhaphuma horsfieldi* White, *Sitophilus zeamais* Motschulsky, *Dendroctonus valens* and *Dastarcus helophoroides*), and even 6 (*T. castaneum*) SNMPs [reported in 59,104]. The role of SNMP1 in pheromone detection has been demonstrated in few Diptera and Lepidoptera species [105–107]. Because of the high degree of conservation of SNMP1 across insect orders, a similar function has been hypothesized also for beetles [105].

The present transcriptomic analysis provides a first understanding of the molecular basis of ladybird olfaction and the role of antennae in male and females *H. axyridis* chemical ecology. Further studies now require functional characterization of the differentially expressed genes we evidenced here to confirm their hypothetical role. Also, transcriptome analysis of other chemosensory tissues such as palps, legs, wings or ovipositor will probably help extending the chemosensory gene repertoire in this species, in concert with genome sequencing.

## Supplementary Materials

Table S1: Sequencing results, Table S1.1: Number of reads obtained by Illumina sequencing, Table S2: GO and InterPro annotations, Table S3: Transcripts per kilobase million (TPM) for the different samples, Table S4: Differential expression analysis, Table S5: Best similarity for odorant receptors (ORs) of *Harmonia axyridis* (Haxy), Table S6: Best similarity for gustatory receptors (GRs) of *Harmonia axyridis* (Haxy), Table S7: Best similarity for ionotropic receptors (IRs) of *Harmonia axyridis* (Haxy), Table S8: Best similarity for odorant-binding proteins (OBPs) of *Harmonia axyridis* (Haxy), Table S9: Best similarity for chemosensory proteins (CSPs) of *Harmonia axyridis* (Haxy), Table S10: Best similarity for sensory neuron membrane proteins (SNMPs) of *Harmonia axyridis* (Haxy), Table S11: Amino acid sequences of *Harmonia axyridis* chemosensory proteins identified in this study, Figure S1. Distribution of *Harmonia axyridis* antennal transcriptome data, Figure S2. Phylogenetic tree of SNMPs.

## Author Contributions

Conceptualization: G.R., E.C., E.J.J.; Methodology: G.R., C.M., N.M., E.C., E.J.J.; Investigation: G.R., A.R., E.J.J; Writing – Original Draft Preparation: G.R., E.C., E.J.J.; Writing – Review & Editing: G.R., N.M., E.C., E.J.J.

## Data Availability

The raw data from Illumina sequencing were deposited in the NCBI Short Read Archive (SRA) database (BioProject ID PRJNA698239).

