## Supplemental tables and figures for "Antennal transcriptome analysis and identification of candidate chemosensory genes of the harlequin ladybird beetle, *Harmonia axyridis* (Pallas) (Coleoptera: Coccinellidae)": Table_S1.1_S1.2_Sequencing.docx

| Sample | *H. axyridis* sex | Reads | Reads>=Q30 |
| --- | --- | --- | --- |
| HA1 | Male | 60,283,120 | 56,142,940 |
| HA3 | Male | 48,988,716 | 44,507,346 |
| HA4 | Male | 61,125,882 | 56,717,050 |
| HA5 | Female | 53,484,392 | 49,795,446 |
| HA6 | Female | 48,838,170 | 45,254,463 |
| HA7 | Female | 60,017,880 | 55,796,899 |
| All statistics are based on contigs of size >= 500 bp, unless otherwise noted (e.g., "# contigs (>= 0 bp)" and "Total length (>= 0 bp)" include all contigs). | | | |

Table S1.1. Number of reads obtained by Illumina sequencing

Table S1.2. Summary of the assembled contigs

| Contigs | | Value |
| --- | --- | --- |
| Total number (>= 500 bp) | | 30,600 |
| # contigs (>= 0 bp) | | 66,728 |
| # contigs (>= 1000 bp) | | 17,208 |
| Largest contig | | 16,954 |
| Total length (>= 500 bp) | | 47,200,406 |
| Total length (>= 0 bp) | | 58,254,799 |
| Total length (>= 1,000 bp) | | 37,810,206 |
| N50 | | 1,981 |
| N75 | | 1,154 |
| L50 | | 7,170 |
| L75 | | 14,968 |
| GC (%) | | 37.82 |
