## Supplemental tables and figures for "Antennal transcriptome analysis and identification of candidate chemosensory genes of the harlequin ladybird beetle, *Harmonia axyridis* (Pallas) (Coleoptera: Coccinellidae)": Table_S4_Differential_expression.docx

| Table S4: Differential expression analysis | | | | |  |  |  |  |
| --- | --- | --- | --- | --- | --- | --- | --- | --- |
| Gene Isoform | log Fold Change (negative is male biased) | P-value | FDR | Accession number | Bit score | E-value | % identity | Description |
| TRINITY_DN14663_c0_g1_i5 | -6.74 | 2.02E-04 | 5.98E-02 | NP_001128425 | 144 | 8.26E-39 | 66.47 | R2D2 protein [Tribolium castaneum]<>PREDICTED: R2D2 protein isoform X1 [Tribolium castaneum]<>PREDICTED: R2D2 protein isoform X1 [Tribolium castaneum]<>hypothetical protein TcasGA2_TC008716 [Tribolium castaneum] |
| TRINITY_DN16216_c0_g1_i4 | -9.69 | 7.78E-06 | 7.70E-03 | KRT86907 | 167 | 1.22E-49 | 69.7 | hypothetical protein AMK59_2413 [Oryctes borbonicus] |
| TRINITY_DN16250_c0_g1_i1 | -6.29 | 2.14E-04 | 6.18E-02 | KYB25350 | 133 | 2.66E-36 | 64.67 | hypothetical protein TcasGA2_TC034890 [Tribolium castaneum] |
| TRINITY_DN16340_c0_g1_i4 | -8.16 | 6.53E-06 | 6.90E-03 | XP_023309869 | 879 | 0.00E+00 | 78.89 | uncharacterized protein LOC108916953 isoform X1 [Anoplophora glabripennis] |
| TRINITY_DN16448_c0_g1_i1 | -7.48 | 3.17E-04 | 7.61E-02 | XP_023016787 | 233 | 1.23E-71 | 62.28 | box C/D snoRNA protein 1 [Leptinotarsa decemlineata] |
| TRINITY_DN16490_c0_g1_i2 | -9.03 | 3.96E-05 | 2.01E-02 | XP_018570025 | 215 | 3.26E-68 | 74.86 | phosphomevalonate kinase [Anoplophora glabripennis] |
| TRINITY_DN16528_c0_g1_i6 | -6.66 | 2.43E-04 | 6.64E-02 | XP_015836321 | 892 | 0.00E+00 | 70.2 | PREDICTED: protein strawberry notch isoform X1 [Tribolium castaneum]<>Protein strawberry notch-like protein [Tribolium castaneum] |
| TRINITY_DN16645_c0_g1_i2 | 7.08 | 4.31E-04 | 9.29E-02 | XP_015838647 | 673 | 0.00E+00 | 88.37 | PREDICTED: LOW QUALITY PROTEIN: formin-2 [Tribolium castaneum] |
| TRINITY_DN17037_c0_g1_i1 | -8.49 | 1.45E-06 | 3.10E-03 | XP_019868615 | 576 | 0.00E+00 | 71.71 | PREDICTED: LOW QUALITY PROTEIN: DNA polymerase iota [Aethina tumida] |
| TRINITY_DN17272_c0_g1_i1 | -7.55 | 4.01E-05 | 2.01E-02 | XP_002098295 | 135 | 1.17E-36 | 64.94 | uncharacterized protein Dyak_GE10305 [Drosophila yakuba]<>similar to Drosophila melanogaster CG18594, partial [Drosophila yakuba]<>uncharacterized protein Dyak_GE10305 [Drosophila yakuba] |
| TRINITY_DN17436_c0_g1_i6 | -8.5 | 1.79E-06 | 3.50E-03 | XP_971282 | 838 | 0.00E+00 | 86.69 | PREDICTED: microprocessor complex subunit DGCR8 [Tribolium castaneum] |
| TRINITY_DN17530_c1_g1_i5 | -8.28 | 2.39E-06 | 4.10E-03 | XP_970935 | 512 | 5.17E-175 | 72.68 | PREDICTED: uncharacterized protein LOC659544 [Tribolium castaneum] |
| TRINITY_DN17814_c0_g1_i4 | -7.8 | 1.25E-05 | 1.02E-02 | XP_008193872 | 282 | 3.66E-90 | 65 | PREDICTED: activating signal cointegrator 1 complex subunit 1 [Tribolium castaneum] |
| TRINITY_DN17894_c0_g1_i1 | 5.42 | 2.93E-04 | 7.29E-02 | XP_023310574 | 228 | 1.39E-73 | 88.81 | cyclic AMP-responsive element-binding protein 1 isoform X9 [Anoplophora glabripennis] |
| TRINITY_DN17942_c0_g1_i2 | -5.99 | 3.44E-05 | 1.91E-02 | ALW95359 | 210 | 1.83E-67 | 82.64 | odorant-binding protein 2 [Cryptolaemus montrouzieri] |
| TRINITY_DN18012_c0_g1_i1 | 7.18 | 3.50E-04 | 8.07E-02 | XP_008191959 | 374 | 4.26E-128 | 78.91 | PREDICTED: serine/threonine-protein phosphatase Pgam5, mitochondrial isoform X2 [Tribolium castaneum] |
| TRINITY_DN18103_c0_g1_i2 | -7.38 | 1.95E-05 | 1.27E-02 | XP_017781058 | 703 | 0.00E+00 | 78.75 | PREDICTED: probable sulfite oxidase, mitochondrial [Nicrophorus vespilloides] |
| TRINITY_DN18236_c0_g1_i1 | -6.89 | 8.88E-05 | 3.63E-02 | XP_966630 | 400 | 4.42E-137 | 88 | PREDICTED: endoplasmic reticulum-Golgi intermediate compartment protein 2 [Tribolium castaneum]<>Endoplasmic reticulum-Golgi intermediate compartment protein 2-like Protein [Tribolium castaneum] |
| TRINITY_DN18263_c0_g1_i2 | 7.44 | 2.92E-04 | 7.29E-02 | KYB25058 | 316 | 3.94E-100 | 78.33 | Coiled-coil domain-containing protein 130 homolog-like Protein [Tribolium castaneum] |
| TRINITY_DN18330_c0_g1_i4 | -6.6 | 1.48E-04 | 5.02E-02 | XP_973618 | 499 | 8.47E-175 | 87.13 | PREDICTED: branched-chain-amino-acid aminotransferase, cytosolic [Tribolium castaneum]<>Branched-chain-amino-acid aminotransferase, cytosolic-like Protein [Tribolium castaneum] |
| TRINITY_DN18590_c0_g1_i1 | -8.02 | 2.21E-04 | 6.26E-02 | XP_022918743 | 158 | 3.26E-40 | 47.4 | zinc finger protein 708-like [Onthophagus taurus] |
| TRINITY_DN18779_c0_g1_i3 | 8.69 | 7.16E-05 | 3.10E-02 | XP_023022327 | 692 | 0.00E+00 | 71 | leucine-rich repeat-containing protein 49 [Leptinotarsa decemlineata] |
| TRINITY_DN18814_c1_g1_i2 | 7.25 | 2.17E-05 | 1.31E-02 | XP_023017977 | 409 | 3.48E-142 | 84.19 | diphthine methyl ester synthase [Leptinotarsa decemlineata] |
| TRINITY_DN18816_c0_g2_i1 | 7.69 | 3.01E-04 | 7.38E-02 | XP_018565165 | 352 | 1.51E-119 | 84.8 | transmembrane protein 41B [Anoplophora glabripennis]<>transmembrane protein 41B-like [Anoplophora glabripennis] |
| TRINITY_DN18877_c0_g1_i4 | -7.54 | 4.31E-04 | 9.29E-02 | XP_018569537 | 543 | 0.00E+00 | 89.38 | probable Dol-P-Man:Man(7)GlcNAc(2)-PP-Dol alpha-1,6-mannosyltransferase [Anoplophora glabripennis] |
| TRINITY_DN18957_c2_g3_i5 | 7.99 | 2.61E-04 | 6.83E-02 | XP_008197345 | 119 | 1.41E-30 | 60.69 | PREDICTED: uncharacterized protein LOC103314117 [Tribolium castaneum]<>COMM domain-containing protein 8-like Protein [Tribolium castaneum] |
| TRINITY_DN19019_c0_g1_i2 | -6.18 | 3.78E-04 | 8.55E-02 | XP_023026692 | 70.1 | 5.09E-11 | 57.14 | odorant receptor 49b-like [Leptinotarsa decemlineata] |
| TRINITY_DN19133_c0_g1_i4 | 7.16 | 1.26E-05 | 1.02E-02 | XP_018578849 | 1453 | 0.00E+00 | 94.31 | transportin-1 [Anoplophora glabripennis]<>transportin-1 [Anoplophora glabripennis] |
| TRINITY_DN19364_c0_g1_i8 | -6.26 | 2.88E-04 | 7.26E-02 | XP_015836731 | 225 | 5.13E-67 | 78.35 | PREDICTED: CREB-regulated transcription coactivator 1 isoform X8 [Tribolium castaneum] |
| TRINITY_DN19450_c0_g1_i1 | 8.11 | 1.38E-04 | 4.77E-02 | XP_018573758 | 541 | 0.00E+00 | 87.95 | UDP-N-acetylglucosamine--dolichyl-phosphate N-acetylglucosaminephosphotransferase [Anoplophora glabripennis] |
| TRINITY_DN19500_c0_g1_i1 | -8.34 | 1.27E-04 | 4.50E-02 | XP_008199002 | 923 | 0.00E+00 | 80.23 | PREDICTED: WASH complex subunit 7 [Tribolium castaneum] |
| TRINITY_DN19597_c0_g2_i2 | 7.86 | 6.58E-06 | 6.90E-03 | XP_018561029 | 575 | 0.00E+00 | 73.25 | transcriptional adapter 2-alpha-like isoform X1 [Anoplophora glabripennis] |
| TRINITY_DN19821_c0_g1_i2 | -8.31 | 6.48E-06 | 6.90E-03 | XP_015835766 | 1051 | 0.00E+00 | 95.06 | PREDICTED: small conductance calcium-activated potassium channel protein isoform X4 [Tribolium castaneum] |
| TRINITY_DN19920_c0_g1_i1 | -8.27 | 1.08E-04 | 4.14E-02 | XP_018567798 | 915 | 0.00E+00 | 77.58 | serine/threonine-protein phosphatase 6 regulatory subunit 3 isoform X1 [Anoplophora glabripennis] |
| TRINITY_DN19987_c1_g1_i8 | -8.8 | 3.72E-05 | 1.97E-02 | XP_019876780 | 697 | 0.00E+00 | 92.71 | PREDICTED: casein kinase II subunit alpha isoform X2 [Aethina tumida] |
| TRINITY_DN20006_c3_g1_i4 | -6.42 | 2.64E-04 | 6.83E-02 | KMQ86558 | 257 | 1.59E-82 | 75.81 | hypothetical protein RF55_14424 [Lasius niger] |
| TRINITY_DN20026_c0_g1_i8 | 8.31 | 7.81E-05 | 3.35E-02 | OCT66771 | 85.1 | 7.85E-17 | 62.5 | hypothetical protein XELAEV_18043022mg [Xenopus laevis] |
| TRINITY_DN20196_c0_g1_i1 | 8.51 | 3.16E-07 | 1.10E-03 | XP_008484571 | 233 | 2.35E-67 | 56.38 | PREDICTED: uncharacterized protein LOC103521244 [Diaphorina citri] |
| TRINITY_DN20196_c0_g1_i5 | 8.51 | 3.13E-07 | 1.10E-03 | KRT78250 | 159 | 1.10E-42 | 80.65 | hypothetical protein AMK59_6683 [Oryctes borbonicus] |
| TRINITY_DN20196_c0_g4_i1 | 7.67 | 2.10E-04 | 6.17E-02 | KMQ87973 | 421 | 4.91E-137 | 71.68 | integrase core domain protein [Lasius niger] |
| TRINITY_DN20417_c1_g2_i1 | 7.93 | 3.33E-04 | 7.89E-02 | APL98297 | 440 | 6.01E-154 | 92.08 | putative DD34D transposase [Bactrocera tryoni] |
| TRINITY_DN20419_c0_g1_i6 | 7.3 | 3.55E-04 | 8.08E-02 | XP_017768046 | 536 | 0.00E+00 | 84.53 | PREDICTED: CLK4-associating serine/arginine rich protein [Nicrophorus vespilloides] |
| TRINITY_DN20419_c0_g1_i7 | -5.86 | 1.25E-04 | 4.50E-02 | XP_017768046 | 533 | 1.76E-180 | 84.53 | PREDICTED: CLK4-associating serine/arginine rich protein [Nicrophorus vespilloides] |
| TRINITY_DN20446_c1_g1_i4 | -7.37 | 3.42E-04 | 7.95E-02 | XP_008196793 | 573 | 0.00E+00 | 89.88 | PREDICTED: heparan sulfate glucosamine 3-O-sulfotransferase 3B1 [Tribolium castaneum] |
| TRINITY_DN20458_c1_g1_i5 | -7.58 | 1.65E-05 | 1.20E-02 | XP_974023 | 679 | 0.00E+00 | 82.93 | PREDICTED: E3 SUMO-protein ligase PIAS2 [Tribolium castaneum]<>E3 SUMO-protein ligase PIAS1-like Protein [Tribolium castaneum] |
| TRINITY_DN20491_c3_g2_i1 | -8.38 | 1.14E-04 | 4.33E-02 | XP_018569695 | 1071 | 0.00E+00 | 74.55 | phosphatidate phosphatase LPIN3 isoform X3 [Anoplophora glabripennis] |
| TRINITY_DN20514_c1_g1_i8 | -8.29 | 1.11E-06 | 2.60E-03 | EFA11165 | 333 | 6.73E-103 | 68.75 | hypothetical protein TcasGA2_TC004772 [Tribolium castaneum] |
| TRINITY_DN20520_c0_g1_i1 | -8.26 | 9.01E-05 | 3.64E-02 | XP_019873670 | 164 | 2.40E-45 | 60.85 | PREDICTED: uncharacterized protein LOC109601822 isoform X2 [Aethina tumida] |
| TRINITY_DN20569_c1_g1_i3 | 7.3 | 4.51E-04 | 9.49E-02 | XP_019556610 | 58.2 | 5.10E-07 | 65.28 | PREDICTED: uncharacterized protein K02A2.6-like [Aedes albopictus] |
| TRINITY_DN20593_c1_g3_i5 | -7.95 | 1.23E-05 | 1.02E-02 | XP_018567680 | 498 | 1.33E-166 | 61.55 | protein tamozhennic [Anoplophora glabripennis] |
| TRINITY_DN20624_c1_g1_i2 | -7.66 | 8.61E-05 | 3.56E-02 | XP_008200998 | 593 | 0.00E+00 | 90.57 | PREDICTED: fruitless isoform X7 [Tribolium castaneum] |
| TRINITY_DN20715_c0_g1_i1 | -7.61 | 1.96E-05 | 1.27E-02 | XP_022902994 | 374 | 2.67E-125 | 76.55 | uncharacterized protein LOC111415502 [Onthophagus taurus] |
| TRINITY_DN20755_c0_g1_i6 | -6.95 | 5.10E-05 | 2.48E-02 | XP_018563470 | 215 | 4.80E-62 | 53.07 | lipase 3-like isoform X2 [Anoplophora glabripennis] |
| TRINITY_DN20798_c2_g1_i2 | -9.76 | 8.77E-13 | 0.00E+00 | BAV13588 | 64.7 | 8.51E-08 | 64.2 | transformer female specific variant [Cyclommatus metallifer finae] |
| TRINITY_DN20828_c0_g1_i3 | -7.96 | 3.59E-06 | 5.30E-03 | XP_008193540 | 497 | 8.07E-164 | 59.04 | PREDICTED: uncharacterized protein LOC103313055 isoform X2 [Tribolium castaneum] |
| TRINITY_DN20879_c0_g2_i2 | -5.29 | 1.92E-04 | 5.83E-02 | XP_021348637 | 68.6 | 1.57E-08 | 55.77 | IgGFc-binding protein-like isoform X1 [Mizuhopecten yessoensis]<>Zonadhesin [Mizuhopecten yessoensis] |
| TRINITY_DN20885_c0_g1_i2 | -7.84 | 5.08E-06 | 6.40E-03 | XP_019865539 | 746 | 0.00E+00 | 75.57 | PREDICTED: serine proteinase stubble [Aethina tumida] |
| TRINITY_DN20987_c0_g1_i7 | -7.89 | 2.13E-09 | 0.00E+00 | XP_970292 | 677 | 0.00E+00 | 96.4 | PREDICTED: tropomodulin isoform X5 [Tribolium castaneum] |
| TRINITY_DN20991_c0_g1_i4 | -7.62 | 6.81E-05 | 2.99E-02 | KYB25718 | 347 | 3.40E-102 | 55.42 | hypothetical protein TcasGA2_TC034125 [Tribolium castaneum] |
| TRINITY_DN21000_c0_g1_i6 | -7.14 | 2.40E-04 | 6.62E-02 | PNF41196 | 461 | 3.40E-144 | 53.28 | hypothetical protein B7P43_G01456 [Cryptotermes secundus]<>hypothetical protein B7P43_G01456 [Cryptotermes secundus] |
| TRINITY_DN21008_c0_g1_i1 | 9.08 | 2.01E-05 | 1.27E-02 | XP_022913568 | 256 | 8.49E-84 | 80 | coiled-coil domain-containing protein 28A isoform X1 [Onthophagus taurus] |
| TRINITY_DN21074_c0_g3_i4 | -7.34 | 3.83E-04 | 8.61E-02 | XP_008194164 | 189 | 1.09E-58 | 72.67 | PREDICTED: phosphatidylinositol N-acetylglucosaminyltransferase subunit P [Tribolium castaneum]<>Phosphatidylinositol N-acetylglucosaminyltransferase subunit P-like Protein [Tribolium castaneum] |
| TRINITY_DN21151_c0_g2_i1 | -9.94 | 1.66E-05 | 1.20E-02 | XP_023026118 | 600 | 0.00E+00 | 82.46 | solute carrier family 35 member F5 [Leptinotarsa decemlineata] |
| TRINITY_DN21177_c0_g1_i1 | -6.58 | 1.51E-04 | 5.05E-02 | XP_022920362 | 256 | 2.78E-77 | 72.64 | A-kinase anchor protein 17A isoform X2 [Onthophagus taurus]<>A-kinase anchor protein 17A isoform X2 [Onthophagus taurus] |
| TRINITY_DN21278_c0_g1_i1 | -7.54 | 2.15E-05 | 1.31E-02 | XP_018331756 | 364 | 7.81E-123 | 71.35 | YJU2 splicing factor homolog [Agrilus planipennis] |
| TRINITY_DN21278_c0_g1_i6 | -7.89 | 2.88E-06 | 4.60E-03 | XP_018331756 | 361 | 9.77E-122 | 94.31 | YJU2 splicing factor homolog [Agrilus planipennis] |
| TRINITY_DN21332_c1_g1_i10 | -8.53 | 9.26E-05 | 3.70E-02 | KYB26781 | 1019 | 0.00E+00 | 88.2 | Phosphatidylinositol 4-phosphate 5-kinase type-1 alpha-like Protein [Tribolium castaneum] |
| TRINITY_DN21345_c0_g1_i1 | -7.48 | 3.14E-04 | 7.61E-02 | XP_008195534 | 1101 | 0.00E+00 | 85.96 | PREDICTED: exocyst complex component 5 [Tribolium castaneum]<>Exocyst complex component 5-like Protein [Tribolium castaneum] |
| TRINITY_DN21444_c0_g1_i3 | -6.56 | 1.85E-04 | 5.72E-02 | XP_018561717 | 493 | 1.71E-172 | 85.62 | WW domain-containing oxidoreductase [Anoplophora glabripennis] |
| TRINITY_DN21462_c0_g2_i4 | -7.9 | 2.34E-04 | 6.56E-02 | XP_017776043 | 961 | 0.00E+00 | 93.14 | PREDICTED: heterogeneous nuclear ribonucleoprotein Q isoform X5 [Nicrophorus vespilloides] |
| TRINITY_DN21515_c0_g1_i4 | 7.75 | 1.55E-04 | 5.09E-02 | XP_023012908 | 436 | 4.84E-142 | 56.62 | ataxin-2-like protein isoform X2 [Leptinotarsa decemlineata] |
| TRINITY_DN21606_c0_g1_i1 | 9.25 | 1.33E-05 | 1.05E-02 | XP_018560907 | 422 | 9.33E-138 | 61.51 | uncharacterized protein LOC108903274 [Anoplophora glabripennis]<>uncharacterized protein LOC108903274 [Anoplophora glabripennis] |
| TRINITY_DN21651_c0_g1_i2 | -6.99 | 3.16E-04 | 7.61E-02 | XP_966551 | 751 | 0.00E+00 | 89.68 | PREDICTED: dnaJ homolog subfamily C member 11 isoform X2 [Tribolium castaneum] |
| TRINITY_DN21749_c0_g5_i3 | -8.55 | 8.16E-07 | 2.10E-03 | XP_023011742 | 553 | 0.00E+00 | 85.2 | homeobox protein extradenticle [Leptinotarsa decemlineata] |
| TRINITY_DN21751_c0_g1_i9 | -6.65 | 8.32E-05 | 3.52E-02 | XP_967662 | 99 | 5.96E-18 | 62.91 | PREDICTED: uncharacterized protein LOC656013 [Tribolium castaneum]<>hypothetical protein TcasGA2_TC008507 [Tribolium castaneum] |
| TRINITY_DN21765_c0_g1_i2 | -10.53 | 3.13E-11 | 0.00E+00 | XP_019873306 | 368 | 2.94E-123 | 69.36 | PREDICTED: LOW QUALITY PROTEIN: inositol polyphosphate 1-phosphatase [Aethina tumida] |
| TRINITY_DN21809_c0_g1_i2 | -8.45 | 1.18E-04 | 4.36E-02 | XP_018336115 | 416 | 1.89E-143 | 90.98 | rRNA 2'-O-methyltransferase fibrillarin [Agrilus planipennis] |
| TRINITY_DN21855_c0_g1_i1 | -8.18 | 1.50E-06 | 3.10E-03 | XP_015840842 | 626 | 0.00E+00 | 85.45 | PREDICTED: endophilin-A isoform X5 [Tribolium castaneum] |
| TRINITY_DN21924_c0_g2_i2 | -7.55 | 1.52E-05 | 1.15E-02 | XP_970128 | 311 | 2.90E-104 | 76.59 | PREDICTED: farnesol dehydrogenase [Tribolium castaneum]<>Dehydrogenase/reductase SDR family protein 7-like [Tribolium castaneum] |
| TRINITY_DN21979_c0_g1_i1 | -3.89 | 3.52E-04 | 8.07E-02 | XP_023014101 | 741 | 0.00E+00 | 92.87 | RNA-binding protein 39 [Leptinotarsa decemlineata] |
| TRINITY_DN22025_c1_g1_i1 | -7.17 | 4.51E-04 | 9.49E-02 | PCG66904 | 239 | 6.79E-73 | 58.59 | hypothetical protein B5V51_7103 [Heliothis virescens] |
| TRINITY_DN22056_c0_g1_i14 | -9 | 5.39E-05 | 2.51E-02 | AJM87404 | 3609 | 0.00E+00 | 91.62 | voltage-sensitive sodium channel [Brassicogethes aeneus] |
| TRINITY_DN22056_c0_g1_i8 | -11.51 | 2.19E-07 | 1.00E-03 | AJM87404 | 3609 | 0.00E+00 | 91.62 | voltage-sensitive sodium channel [Brassicogethes aeneus] |
| TRINITY_DN22056_c0_g1_i9 | -9.75 | 9.95E-06 | 8.80E-03 | AJM87404 | 3655 | 0.00E+00 | 92.48 | voltage-sensitive sodium channel [Brassicogethes aeneus] |
| TRINITY_DN22141_c0_g1_i3 | -3.08 | 1.78E-05 | 1.24E-02 | XP_015834462 | 1810 | 0.00E+00 | 85.01 | PREDICTED: uncharacterized protein LOC657912 isoform X2 [Tribolium castaneum] |
| TRINITY_DN22175_c0_g2_i2 | -9.6 | 4.99E-08 | 4.00E-04 | XP_022918167 | 622 | 0.00E+00 | 75.68 | protein bric-a-brac 1-like isoform X1 [Onthophagus taurus]<>protein bric-a-brac 1-like isoform X1 [Onthophagus taurus] |
| TRINITY_DN22175_c0_g2_i4 | -9.91 | 4.44E-06 | 6.10E-03 | XP_022918167 | 622 | 0.00E+00 | 75.68 | protein bric-a-brac 1-like isoform X1 [Onthophagus taurus]<>protein bric-a-brac 1-like isoform X1 [Onthophagus taurus] |
| TRINITY_DN22297_c0_g1_i1 | -5.75 | 1.62E-04 | 5.19E-02 | XP_018575393 | 680 | 0.00E+00 | 60.78 | integrin beta-PS-like [Anoplophora glabripennis] |
| TRINITY_DN22399_c0_g2_i1 | -7.47 | 1.40E-05 | 1.08E-02 | XP_008194992 | 1032 | 0.00E+00 | 95.64 | PREDICTED: segment polarity protein dishevelled homolog DVL-3 isoform X2 [Tribolium castaneum] |
| TRINITY_DN22410_c0_g1_i3 | -9.61 | 3.92E-04 | 8.77E-02 | APR62727 | 3700 | 0.00E+00 | 99.72 | vitellogenin 1 [Harmonia axyridis] |
| TRINITY_DN22410_c0_g2_i1 | -7.96 | 8.73E-06 | 8.20E-03 | APR62728 | 3706 | 0.00E+00 | 99.61 | vitellogenin 2 [Harmonia axyridis] |
| TRINITY_DN22446_c0_g1_i4 | -6.82 | 1.28E-04 | 4.50E-02 | AVM18964 | 157 | 2.59E-47 | 87.63 | odorant binding protein 38, partial [Holotrichia parallela] |
| TRINITY_DN22453_c0_g1_i1 | -8.24 | 2.22E-04 | 6.26E-02 | XP_966456 | 348 | 2.62E-114 | 75.96 | PREDICTED: probable glutamate--tRNA ligase, mitochondrial [Tribolium castaneum]<>putative glutamate--tRNA ligase, mitochondrial-like Protein [Tribolium castaneum] |
| TRINITY_DN22469_c0_g1_i5 | 7.02 | 7.17E-06 | 7.30E-03 | XP_019762299 | 189 | 1.01E-53 | 84.4 | PREDICTED: uncharacterized protein LOC109539132 [Dendroctonus ponderosae] |
| TRINITY_DN22525_c0_g2_i7 | 9.12 | 3.38E-05 | 1.91E-02 | XP_968830 | 1185 | 0.00E+00 | 68.58 | PREDICTED: myotubularin-related protein 3 isoform X2 [Tribolium castaneum]<>Myotubularin-related protein 3-like Protein [Tribolium castaneum] |
| TRINITY_DN22528_c0_g2_i1 | -6.69 | 4.62E-04 | 9.61E-02 | XP_015839312 | 763 | 0.00E+00 | 93.26 | PREDICTED: echinoderm microtubule-associated protein-like CG42247 isoform X1 [Tribolium castaneum] |
| TRINITY_DN22599_c0_g2_i6 | -6.09 | 1.90E-05 | 1.27E-02 | XP_002046155 | 699 | 0.00E+00 | 71.75 | uncharacterized protein Dvir_GJ12672, isoform I [Drosophila virilis]<>uncharacterized protein Dvir_GJ12672, isoform I [Drosophila virilis] |
| TRINITY_DN22623_c0_g1_i2 | -6.21 | 5.73E-06 | 6.80E-03 | OLP75034 | 104 | 4.37E-21 | 63.03 | hypothetical protein AK812_SmicGene45246 [Symbiodinium microadriaticum] |
| TRINITY_DN22638_c0_g1_i3 | -9.21 | 3.45E-05 | 1.91E-02 | XP_018571016 | 1081 | 0.00E+00 | 74.73 | transmembrane protein 245 [Anoplophora glabripennis] |
| TRINITY_DN22638_c0_g1_i4 | -9.58 | 8.14E-06 | 7.80E-03 | XP_018571016 | 1081 | 0.00E+00 | 74.73 | transmembrane protein 245 [Anoplophora glabripennis] |
| TRINITY_DN22643_c0_g1_i1 | 8.76 | 4.55E-07 | 1.50E-03 | XP_008191255 | 362 | 9.35E-109 | 61.26 | PREDICTED: uncharacterized protein LOC659080 [Tribolium castaneum]<>PREDICTED: uncharacterized protein LOC659080 [Tribolium castaneum] |
| TRINITY_DN22734_c0_g1_i3 | -8.58 | 6.63E-05 | 2.94E-02 | KYB27443 | 809 | 0.00E+00 | 73.41 | Tubulin glycylase 3A-like Protein [Tribolium castaneum] |
| TRINITY_DN22734_c0_g1_i7 | 4.6 | 4.56E-04 | 9.53E-02 | KYB27443 | 810 | 0.00E+00 | 73.3 | Tubulin glycylase 3A-like Protein [Tribolium castaneum] |
| TRINITY_DN22794_c0_g3_i2 | -6.6 | 1.40E-04 | 4.80E-02 | XP_015838904 | 659 | 0.00E+00 | 95.36 | PREDICTED: Down syndrome cell adhesion molecule isoform X32 [Tribolium castaneum] |
| TRINITY_DN22796_c0_g1_i6 | -9.67 | 1.95E-06 | 3.50E-03 | XP_018579331 | 1916 | 0.00E+00 | 80.33 | rho guanine nucleotide exchange factor 18 isoform X4 [Anoplophora glabripennis] |
| TRINITY_DN22814_c0_g1_i1 | -8.63 | 5.43E-05 | 2.51E-02 | XP_018579710 | 1274 | 0.00E+00 | 81.81 | nuclear export mediator factor NEMF homolog [Anoplophora glabripennis] |
| TRINITY_DN23052_c0_g1_i3 | -7.98 | 2.65E-04 | 6.83E-02 | XP_019872456 | 1231 | 0.00E+00 | 71.64 | PREDICTED: protein timeless-like [Aethina tumida] |
| TRINITY_DN23152_c1_g1_i1 | -5.67 | 1.17E-04 | 4.36E-02 | XP_015838785 | 1941 | 0.00E+00 | 92.31 | PREDICTED: Ca(2+)/calmodulin-responsive adenylate cyclase isoform X8 [Tribolium castaneum] |
| TRINITY_DN23188_c0_g2_i4 | -7.17 | 4.39E-04 | 9.40E-02 | AQS83399 | 338 | 5.01E-109 | 83.54 | boule [Diabrotica virgifera] |
| TRINITY_DN23188_c0_g2_i8 | 7.09 | 4.02E-04 | 8.87E-02 | AQS83399 | 397 | 8.81E-131 | 76.51 | boule [Diabrotica virgifera] |
| TRINITY_DN23206_c0_g1_i4 | -6.39 | 1.98E-04 | 5.91E-02 | XP_018562050 | 1718 | 0.00E+00 | 98.62 | sodium/potassium-transporting ATPase subunit alpha isoform X2 [Anoplophora glabripennis] |
| TRINITY_DN23220_c0_g1_i5 | -6.57 | 2.21E-05 | 1.31E-02 | XP_019866742 | 482 | 8.15E-165 | 90 | PREDICTED: LOW QUALITY PROTEIN: calcium-binding mitochondrial carrier protein Aralar1 [Aethina tumida] |
| TRINITY_DN23234_c0_g1_i4 | 9.45 | 1.13E-05 | 9.80E-03 | XP_975506 | 214 | 2.65E-59 | 53.66 | PREDICTED: double-stranded RNA-binding protein Staufen homolog 2 [Tribolium castaneum]<>Maternal effect protein staufen-like Protein [Tribolium castaneum] |
| TRINITY_DN23377_c0_g1_i4 | -8.56 | 2.96E-06 | 4.60E-03 | XP_019873410 | 573 | 0.00E+00 | 73.71 | PREDICTED: UNC93-like protein isoform X1 [Aethina tumida] |
| TRINITY_DN23483_c0_g1_i8 | -9.43 | 3.50E-05 | 1.91E-02 | XP_967114 | 884 | 0.00E+00 | 79.85 | PREDICTED: protein fem-1 homolog CG6966 [Tribolium castaneum]<>Protein fem-1 homolog CG6966-like Protein [Tribolium castaneum] |
| TRINITY_DN23568_c0_g1_i2 | -8.43 | 1.26E-04 | 4.50E-02 | XP_015836282 | 1648 | 0.00E+00 | 76.38 | PREDICTED: activating signal cointegrator 1 complex subunit 3 [Tribolium castaneum]<>Putative U5 small nuclear ribonucleoprotein 200 kDa helicase-like protein [Tribolium castaneum] |
| TRINITY_DN23588_c0_g1_i3 | -9.43 | 2.09E-07 | 1.00E-03 | XP_023029951 | 1828 | 0.00E+00 | 92.02 | vinculin isoform X1 [Leptinotarsa decemlineata] |
| TRINITY_DN23753_c0_g1_i6 | -8.16 | 1.64E-04 | 5.20E-02 | XP_018569640 | 603 | 0.00E+00 | 65.86 | putative tyrosine-protein kinase Wsck [Anoplophora glabripennis] |
| TRINITY_DN23927_c0_g1_i1 | -7.35 | 1.74E-05 | 1.23E-02 | XP_023311314 | 711 | 0.00E+00 | 91.02 | uncharacterized protein CG3556 isoform X2 [Anoplophora glabripennis] |
| TRINITY_DN24018_c0_g1_i4 | -8.59 | 5.21E-05 | 2.50E-02 | XP_018327190 | 989 | 0.00E+00 | 57.7 | PHD finger protein rhinoceros-like [Agrilus planipennis] |
| TRINITY_DN24107_c0_g1_i1 | -8.57 | 5.36E-07 | 1.60E-03 | XP_018576475 | 450 | 4.16E-146 | 91.77 | tetratricopeptide repeat protein 14 homolog [Anoplophora glabripennis] |
| TRINITY_DN24164_c0_g3_i3 | 4.95 | 3.99E-04 | 8.86E-02 | XP_019880464 | 566 | 0.00E+00 | 98.21 | PREDICTED: A disintegrin and metalloproteinase with thrombospondin motifs 16 [Aethina tumida] |
| TRINITY_DN24275_c0_g1_i4 | -7.34 | 2.54E-04 | 6.77E-02 | BAA75470 | 228 | 3.82E-68 | 84.71 | prophenoloxidase [Tenebrio molitor] |
| TRINITY_DN24320_c0_g1_i5 | -8.85 | 1.23E-04 | 4.50E-02 | ACL50549 | 860 | 0.00E+00 | 99.52 | trehalase-2 [Harmonia axyridis] |
| TRINITY_DN24335_c0_g1_i3 | -8.97 | 4.00E-05 | 2.01E-02 | XP_001809587 | 808 | 0.00E+00 | 69.77 | PREDICTED: zinc finger protein 2 homolog [Tribolium castaneum] |
| TRINITY_DN24382_c1_g1_i1 | -7.53 | 2.43E-05 | 1.42E-02 | XP_025837031 | 97.8 | 7.98E-21 | 56.85 | glucose dehydrogenase [FAD, quinone]-like [Agrilus planipennis] |
| TRINITY_DN24692_c1_g2_i1 | -6.41 | 2.18E-04 | 6.24E-02 | XP_017772831 | 162 | 1.23E-44 | 82.57 | PREDICTED: uncharacterized protein LOC108559950 isoform X2 [Nicrophorus vespilloides] |
| TRINITY_DN24782_c0_g1_i5 | 7.31 | 1.16E-07 | 8.00E-04 | APP94029 | 784 | 0.00E+00 | 94.58 | shaggy [Colaphellus bowringi] |
| TRINITY_DN24800_c1_g1_i3 | -7.65 | 4.42E-04 | 9.40E-02 | XP_017771289 | 167 | 3.19E-40 | 60.38 | PREDICTED: RB1-inducible coiled-coil protein 1-like isoform X2 [Nicrophorus vespilloides] |
| TRINITY_DN25107_c0_g1_i3 | 10.96 | 4.61E-06 | 6.10E-03 | XP_968598 | 1578 | 0.00E+00 | 88.03 | PREDICTED: transient-receptor-potential-like protein isoform X1 [Tribolium castaneum]<>TRP gamma [Tribolium castaneum] |
| TRINITY_DN25157_c0_g2_i2 | -8.48 | 1.02E-04 | 3.98E-02 | KYB28281 | 2131 | 0.00E+00 | 94.34 | hypothetical protein TcasGA2_TC034635 [Tribolium castaneum] |
| TRINITY_DN25275_c15_g1_i15 | -7.13 | 2.15E-05 | 1.31E-02 | APL98293 | 278 | 3.26E-91 | 93.51 | putative DD34D transposase [Bactrocera tryoni] |
| TRINITY_DN25275_c15_g1_i16 | 7.36 | 2.81E-04 | 7.14E-02 | AAA28265 | 217 | 2.31E-67 | 90.32 | mariner transposase [Chrysoperla plorabunda]<>mariner transposase [Chrysoperla plorabunda] |
