## Supplemental tables and figures for "Antennal transcriptome analysis and identification of candidate chemosensory genes of the harlequin ladybird beetle, *Harmonia axyridis* (Pallas) (Coleoptera: Coccinellidae)": Table_S5_ORs.docx

| Table S5: Best similarity for odorant receptors (ORs) of *Harmonia axyridis* (Haxy) | | | | | | | |  |  |
| --- | --- | --- | --- | --- | --- | --- | --- | --- | --- |
| Name | ID | length (aa) | ORF status | TMHMM | Bit score | E-value | % identity | Accession number | Description |
| HaxyOrco | TRINITY_DN24509_c0_g1::TRINITY_DN24509_c0_g1_i2 | 480 | complete | 7 | 786.0 | 0 | 79 | XP_008194693 | PREDICTED: odorant receptor coreceptor [Tribolium castaneum] |
| HaxyOR2 | TRINITY_DN16089_c0_g1::TRINITY_DN16089_c0_g1_i2 | 337 | 5prime_partial | 5 | 143.0 | 5.12e-36 | 33 | EFA09299 | odorant receptor 110 [Tribolium castaneum] |
| HaxyOR3 | TRINITY_DN18065_c0_g1::TRINITY_DN18065_c0_g1_i2 | 374 | complete | 6 | 130.0 | 1.04e-30 | 27 | XP_015838469 | PREDICTED: uncharacterized protein LOC107398561 [Tribolium castaneum] |
| HaxyOR4 | TRINITY_DN18831_c0_g1::TRINITY_DN18831_c0_g1_i1 | 395 | complete | 4 | 116.0 | 2.32e-25 | 31 | XP_023026692 | odorant receptor 49b-like [Leptinotarsa decemlineata] |
| HaxyOR5 | TRINITY_DN19019_c0_g1::TRINITY_DN19019_c0_g1_i1 | 316 | 5prime_partial | 6 | 75.5 | 1.51e-11 | 24 | XP_023309827 | odorant receptor Or2-like [Anoplophora glabripennis] |
| HaxyOR6 | TRINITY_DN19257_c0_g1::TRINITY_DN19257_c0_g1_i2 | 367 | complete | 6 | 135.0 | 1.5e-32 | 25 | EFA09299 | odorant receptor 110 [Tribolium castaneum] |
| HaxyOR7 | TRINITY_DN19287_c0_g1::TRINITY_DN19287_c0_g1_i2 | 397 | 5prime_partial | 5 | 243.0 | 1.56e-73 | 38 | AJO62235 | olfactory receptor OR16 |
| HaxyOR8 | TRINITY_DN19317_c0_g1::TRINITY_DN19317_c0_g1_i1 | 412 | complete | 5 | 107.0 | 3.46e-22 | 24 | XP_019874691 | PREDICTED: odorant receptor 67c-like [Aethina tumida] |
| HaxyOR9 | TRINITY_DN19681_c0_g1::TRINITY_DN19681_c0_g1_i1 | 382 | 5prime_partial | 6 | 229.0 | 4.32e-68 | 33 | XP_015833042 | PREDICTED: odorant receptor Or1 isoform X4 [Tribolium castaneum] |
| HaxyOR10 | TRINITY_DN19682_c4_g1::TRINITY_DN19682_c4_g1_i1 | 388 | complete | 6 | 75.5 | 2.6e-11 | 24 | EFA04742 | odorant receptor 331 [Tribolium castaneum] |
| HaxyOR11 | TRINITY_DN20080_c2_g1::TRINITY_DN20080_c2_g1_i1 | 339 | complete | 5 | 62.0 | 7e-07 | 24 | EEZ97786.1 | odorant receptor 309 [Tribolium castaneum] |
| HaxyOR12 | TRINITY_DN20680_c0_g1::TRINITY_DN20680_c0_g1_i1 | 409 | complete | 6 | 81.6 | 2.57e-13 | 24 | XP_019755730 | PREDICTED: odorant receptor 49b-like isoform X1 [Dendroctonus ponderosae] |
| HaxyOR13 | TRINITY_DN21300_c2_g1::TRINITY_DN21300_c2_g1_i1 | 398 | 5prime_partial | 4 | 227.0 | 5.88e-67 | 32 | XP_015840910 | PREDICTED: odorant receptor Or1-like isoform X1 [Tribolium castaneum] |
| HaxyOR14 | TRINITY_DN21390_c0_g2::TRINITY_DN21390_c0_g2_i1 | 399 | complete | 6 | 125.0 | 1.18e-28 | 28 | XP_023026692 | odorant receptor 49b-like [Leptinotarsa decemlineata] |
| HaxyOR15 | TRINITY_DN21643_c0_g1::TRINITY_DN21643_c0_g1_i2 | 266 | internal | 4 | 127.0 | 1.67e-29 | 31 | XP_015839882 | PREDICTED: uncharacterized protein LOC663463 [Tribolium castaneum] |
| HaxyOR16 | TRINITY_DN21983_c0_g1::TRINITY_DN21983_c0_g1_i2 | 398 | 5prime_partial | 6 | 114.0 | 2.47e-24 | 23 | XP_023026692 | odorant receptor 49b-like [Leptinotarsa decemlineata] |
| HaxyOR17 | TRINITY_DN22213_c4_g3::TRINITY_DN22213_c4_g3_i1 | 410 | 5prime_partial | 4 | 212.0 | 2.68e-61 | 31 | AJO62225 | olfactory receptor OR6 |
| HaxyOR18 | TRINITY_DN22336_c0_g1::TRINITY_DN22336_c0_g1_i2 | 397 | complete | 6 | 69.7 | 5e-09 | 27 | ALR72565.1 | odorant receptor OR20 [Colaphellus bowringi] |
| HaxyOR19 | TRINITY_DN22419_c0_g1::TRINITY_DN22419_c0_g1_i1 | 222 | 3prime_partial | 4 | 92.4 | 3.98e-19 | 27 | XP_008197157 | PREDICTED: odorant receptor Or1-like [Tribolium castaneum] |
| HaxyOR20 | TRINITY_DN22684_c0_g1::TRINITY_DN22684_c0_g1_i3 | 394 | complete | 7 | 241.0 | 1.01e-72 | 32 | XP_015833042 | PREDICTED: odorant receptor Or1 isoform X4 [Tribolium castaneum] |
| HaxyOR21 | TRINITY_DN22897_c0_g1::TRINITY_DN22897_c0_g1_i2 | 163 | 5prime_partial | 2 | 56.2 | 2.85e-06 | 31 | EFA02946 | odorant receptor 159 [Tribolium castaneum] |
| HaxyOR22 | TRINITY_DN23027_c0_g1::TRINITY_DN23027_c0_g1_i4 | 165 | 5prime_partial | 2 | 93.6 | 2.58e-19 | 32 | XP_023026692 | odorant receptor 49b-like [Leptinotarsa decemlineata] |
| HaxyOR23 | TRINITY_DN23903_c0_g1::TRINITY_DN23903_c0_g1_i1 | 408 | 5prime_partial | 6 | 75.9 | 2.38e-11 | 25 | NP_001166620 | olfactory receptor 63 [Bombyx mori]<>olfactory receptor [Bombyx mori] |
| HaxyOR24 | TRINITY_DN137_c0_g1::TRINITY_DN137_c0_g1_i1 | 142 | 5prime_partial | 0 | 57.4 | 7.5e-07 | 30 | XP_022904516 | odorant receptor 85b-like [Onthophagus taurus] |
| HaxyOR25 | TRINITY_DN29248_c0_g1::TRINITY_DN29248_c0_g1_i1 | 102 | internal | 2 | 58.9 | 5.52e-08 | 31 | EFA01395 | odorant receptor 185 [Tribolium castaneum] |
| HaxyOR26 | TRINITY_DN5091_c0_g1::TRINITY_DN5091_c0_g1_i2 | 276 | internal | 4 | 85.1 | 3.32e-15 | 24 | EFA02958 | odorant receptor 315 [Tribolium castaneum] |
