## Supplemental tables and figures for "Antennal transcriptome analysis and identification of candidate chemosensory genes of the harlequin ladybird beetle, *Harmonia axyridis* (Pallas) (Coleoptera: Coccinellidae)": Table_S6_GRs.docx

| Table S6: Best similarity for gustatory receptors (GRs) of *Harmonia axyridis* (Haxy) | | | | | | | | |  |
| --- | --- | --- | --- | --- | --- | --- | --- | --- | --- |
| Name | ID | length (aa) | ORF status | TMHMM | Bit score | E-value | % identity | Accession number | Description |
| HaxyGR1 | TRINITY_DN14734_c0_g1::TRINITY_DN14734_c0_g1_i1 | 210 | internal | 4 | 150.0 | 6.37e-40 | 39.901 | EFA04718 | gustatory receptor 12 [Tribolium castaneum] |
| HaxyGR2 | TRINITY_DN14855_c0_g1::TRINITY_DN14855_c0_g1_i1 | 440 | complete | 7 | 232.0 | 3.88e-68 | 33.882 | AVN97874 | gustatory receptor 9 |
| HaxyGR3 | TRINITY_DN15534_c0_g1::TRINITY_DN15534_c0_g1_i1 | 448 | 5prime_partial | 7 | 289.0 | 3.39e-90 | 37.337 | XP_015836176 | PREDICTED: gustatory receptor for sugar taste 64e-like isoform X2 [Tribolium castaneum]<>gustatory receptor candidate 29 [Tribolium castaneum]<>gustatory receptor candidate 7 [Tribolium castaneum] |
| HaxyGR4 | TRINITY_DN17179_c0_g1::TRINITY_DN17179_c0_g1_i3 | 361 | 5prime_partial | 6 | 191.0 | 1.97e-53 | 29.050 | XP_015836176 | PREDICTED: gustatory receptor for sugar taste 64e-like isoform X2 [Tribolium castaneum]<>gustatory receptor candidate 29 [Tribolium castaneum]<>gustatory receptor candidate 7 [Tribolium castaneum] |
| HaxyGR5 | TRINITY_DN18813_c0_g1::TRINITY_DN18813_c0_g1_i3 | 153 | 5prime_partial | 0 | 100.0 | 4.25e-24 | 41.958 | XP_023030044 | uncharacterized protein LOC111517976 [Leptinotarsa decemlineata] |
| HaxyGR6 | TRINITY_DN19843_c0_g1::TRINITY_DN19843_c0_g1_i1 | 381 | internal | 7 | 179.0 | 8.39e-49 | 32.514 | XP_015836206 | PREDICTED: gustatory receptor for sugar taste 43a [Tribolium castaneum] |
| HaxyGR7 | TRINITY_DN20841_c0_g1::TRINITY_DN20841_c0_g1_i1 | 284 | 3prime_partial | 4 | 137.0 | 3.3e-33 | 31.273 | KYB27391 | Gustatory receptor for sugar taste 64e-like Protein [Tribolium castaneum] |
| HaxyGR8 | TRINITY_DN20841_c0_g2::TRINITY_DN20841_c0_g2_i1 | 122 | internal | 1 | 111.0 | 1.1e-26 | 49.580 | XP_019760210 | PREDICTED: gustatory receptor for sugar taste 64f-like [Dendroctonus ponderosae] |
| HaxyGR9 | TRINITY_DN21679_c0_g1::TRINITY_DN21679_c0_g1_i1 | 392 | complete | 8 | 351.0 | 2.41e-115 | 45.758 | XP_008193165 | PREDICTED: gustatory receptor Gr83 isoform X2 [Tribolium castaneum] |
| HaxyGR10 | TRINITY_DN21760_c0_g2::TRINITY_DN21760_c0_g2_i1 | 114 | 5prime_partial | 1 | 57.8 | 1.99e-08 | 37.209 | XP_018569254 | gustatory receptor 68a-like [Anoplophora glabripennis] |
| HaxyGR11 | TRINITY_DN22663_c0_g1::TRINITY_DN22663_c0_g1_i1 | 196 | internal | 3 | 343.0 | 5.49e-115 | 80.513 | XP_022903591 | gustatory and odorant receptor 22-like [Onthophagus taurus] |
| HaxyGR12 | TRINITY_DN22663_c0_g2::TRINITY_DN22663_c0_g2_i1 | 117 | 3prime_partial | 1 | 167.0 | 1.03e-47 | 68.644 | XP_018563765 | gustatory and odorant receptor 22-like [Anoplophora glabripennis] |
| HaxyGR13 | TRINITY_DN22663_c0_g3::TRINITY_DN22663_c0_g3_i1 | 122 | 5prime_partial | 0 | 180.0 | 4.49e-53 | 70.940 | NP_001161916 | gustatory receptor 2 [Tribolium castaneum]<>gustatory receptor candidate 47 [Tribolium castaneum]<>gustatory receptor candidate 26 [Tribolium castaneum]<>gustatory receptor candidate 6 [Tribolium castaneum] |
| HaxyGR14 | TRINITY_DN24512_c0_g1::TRINITY_DN24512_c0_g1_i1 | 136 | internal | 1 | 116.0 | 1.71e-28 | 41.912 | XP_008195245 | PREDICTED: putative gustatory receptor 28a [Tribolium castaneum] |
| HaxyGR15 | TRINITY_DN6575_c0_g1::TRINITY_DN6575_c0_g1_i1 | 137 | internal | 2 | 237.0 | 3.52e-74 | 81.752 | XP_017774851 | PREDICTED: gustatory and odorant receptor 24 [Nicrophorus vespilloides] |
| HaxyGR16 | TRINITY_DN24470_c1_g1::TRINITY_DN24470_c1_g1_i1 | 165 | 5prime_partial | 3 | 43.9 | 0.069 | 31 | XP_023023560.1 | odorant receptor 47b-like [Leptinotarsa decemlineata] |
| HaxyGR17 | TRINITY_DN25825_c0_g1::TRINITY_DN25825_c0_g1_i1 | 104 | internal | 0 | 50.4 | 7e-05 | 30 | EFA02941.1 | odorant receptor 93 [Tribolium castaneum] |
