## Supplemental tables and figures for "Antennal transcriptome analysis and identification of candidate chemosensory genes of the harlequin ladybird beetle, *Harmonia axyridis* (Pallas) (Coleoptera: Coccinellidae)": Table_S7_IRs.docx

| Table S7: Best similarity for ionotropic receptors (IRs) of *Harmonia axyridis* (Haxy) | | | | | | |  |  |  |
| --- | --- | --- | --- | --- | --- | --- | --- | --- | --- |
| Name | ID | length (aa) | ORF status | TMHMM | Bit score | E-value | % identity | Accession number | Description |
| HaxyIR25a.7 | TRINITY_DN1065_c0_g1::TRINITY_DN1065_c0_g1_i1 | 117 | internal | 0 | 134.0 | 6.26e-34 | 53.636 | XP_002066605 | ionotropic receptor 25a [Drosophila willistoni] |
| HaxyIR40a.1 | TRINITY_DN11365_c0_g1::TRINITY_DN11365_c0_g1_i1 | 127 | internal | 0 | 183.0 | 2.09e-56 | 70.079 | AJO62241 | chemosensory ionotropic receptor IR3 |
| HaxyIR1 | TRINITY_DN1360_c0_g1::TRINITY_DN1360_c0_g1_i1 | 223 | internal | 0 | 165.0 | 2.33e-43 | 37.900 | XP_023022175 | ionotropic receptor 25a [Leptinotarsa decemlineata] |
| HaxyIR2 | TRINITY_DN15513_c0_g1::TRINITY_DN15513_c0_g1_i1 | 201 | internal | 0 | 73.6 | 9.11e-12 | 25.658 | PNF21465 | hypothetical protein B7P43_G13524 [Cryptotermes secundus] |
| HaxyIR56e.1 | TRINITY_DN16178_c0_g1::TRINITY_DN16178_c0_g1_i1 | 301 | complete | 7 | 340.0 | 7.52e-114 | 56.587 | ETN66287 | nmda receptor glutamate-binding chain [Anopheles darlingi] |
| HaxyIR8a.1 | TRINITY_DN16593_c0_g1::TRINITY_DN16593_c0_g1_i1 | 184 | internal | 0 | 168.0 | 5.48e-45 | 44.086 | ALR72538 | ionotropic receptor 8a [Colaphellus bowringi] |
| HaxyIR8a.2 | TRINITY_DN16816_c0_g1::TRINITY_DN16816_c0_g1_i2 | 122 | internal | 0 | 188.0 | 2.63e-53 | 65.041 | XP_023311227 | ionotropic receptor 25a [Anoplophora glabripennis] |
| HaxyIR75q.1 | TRINITY_DN16963_c0_g1::TRINITY_DN16963_c0_g1_i11 | 236 | internal | 2 | 236.0 | 7.39e-72 | 47.458 | AVH87301 | ionotropic receptor 13 [Holotrichia parallela] |
| HaxyIR68a.1 | TRINITY_DN16973_c0_g1::TRINITY_DN16973_c0_g1_i1 | 268 | 5prime_partial | 2 | 381.0 | 4.93e-125 | 66.790 | XP_015839172 | PREDICTED: glutamate receptor ionotropic kainate 5 [Tribolium castaneum] |
| HaxyIR25a.1 | TRINITY_DN17144_c0_g3::TRINITY_DN17144_c0_g3_i1 | 179 | 5prime_partial | 1 | 173.0 | 8.34e-47 | 48.387 | AGJ51188 | olfactory ionotropic receptor IR25a [Panulirus argus] |
| HaxyIR3 | TRINITY_DN19744_c0_g1::TRINITY_DN19744_c0_g1_i1 | 689 | internal | 4 | 217.0 | 3e-58 | 28.699 | EFA08518 | hypothetical protein TcasGA2_TC006171 [Tribolium castaneum] |
| HaxyIR64a.1 | TRINITY_DN22273_c0_g1::TRINITY_DN22273_c0_g1_i1 | 643 | complete | 4 | 629.0 | 0 | 53.729 | XP_008196049 | PREDICTED: glutamate receptor ionotropic |
| HaxyIR4 | TRINITY_DN22389_c0_g1::TRINITY_DN22389_c0_g1_i3 | 563 | 5prime_partial | 4 | 149.0 | 1.63e-35 | 26.895 | XP_017774907 | PREDICTED: uncharacterized protein LOC108561472 [Nicrophorus vespilloides] |
| HaxyIR21a | TRINITY_DN22807_c1_g2::TRINITY_DN22807_c1_g2_i2 | 568 | internal | 2 | 614.0 | 0 | 55.153 | XP_019864916 | PREDICTED: ionotropic receptor 21a [Aethina tumida] |
| HaxyIR75q.2 | TRINITY_DN23506_c0_g1::TRINITY_DN23506_c0_g1_i1 | 644 | 5prime_partial | 4 | 438.0 | 5.63e-143 | 40.824 | APC94348 | ionotropic receptor 5 |
| HaxyIR76b | TRINITY_DN24055_c0_g1::TRINITY_DN24055_c0_g1_i1 | 541 | complete | 4 | 554.0 | 0 | 51.601 | AJO62243 | chemosensory ionotropic receptor IR5 [Tenebrio molitor] |
| HaxyIR25a.2 | TRINITY_DN24365_c2_g2::TRINITY_DN24365_c2_g2_i1 | 917 | complete | 3 | 1147.0 | 0 | 59.259 | AJO62244 | chemosensory ionotropic receptor IR6 [Tenebrio molitor] |
| HaxyIR75s | TRINITY_DN24542_c1_g1::TRINITY_DN24542_c1_g1_i1 | 160 | internal | 0 | 119.0 | 2.37e-29 | 45.926 | AVN97883 | ionotropic receptor 1 |
| HaxyIR75q.3 | TRINITY_DN24777_c4_g1::TRINITY_DN24777_c4_g1_i1 | 607 | 5prime_partial | 3 | 416.0 | 5.92e-135 | 37.990 | APC94348 | ionotropic receptor 5 |
| HaxyIR93a | TRINITY_DN24818_c1_g1::TRINITY_DN24818_c1_g1_i1 | 628 | internal | 2 | 546.0 | 0 | 44.928 | XP_018576792 | ionotropic receptor 93a isoform X1 [Anoplophora glabripennis] |
| HaxyIR25a.3 | TRINITY_DN2489_c0_g1::TRINITY_DN2489_c0_g1_i1 | 111 | internal | 0 | 144.0 | 1.09e-37 | 58.036 | AJO62244 | chemosensory ionotropic receptor IR6 [Tenebrio molitor] |
| HaxyIR8a.3 | TRINITY_DN25068_c4_g1::TRINITY_DN25068_c4_g1_i1 | 915 | 5prime_partial | 4 | 1095.0 | 0 | 60.090 | XP_968346 | PREDICTED: glutamate receptor ionotropic |
| HaxyIR25a.4 | TRINITY_DN25200_c1_g1::TRINITY_DN25200_c1_g1_i1 | 938 | complete | 3 | 1394.0 | 0 | 75.200 | AJO62244 | chemosensory ionotropic receptor IR6 [Tenebrio molitor] |
| HaxyIR25a.5 | TRINITY_DN28320_c0_g1::TRINITY_DN28320_c0_g1_i1 | 212 | internal | 2 | 347.0 | 7.43e-111 | 74.057 | XP_017786976 | PREDICTED: ionotropic receptor 25a [Nicrophorus vespilloides] |
| HaxyIR25a.6 | TRINITY_DN5298_c0_g1::TRINITY_DN5298_c0_g1_i1 | 150 | 5prime_partial | 1 | 219.0 | 5.7e-64 | 73.381 | AJO62244 | chemosensory ionotropic receptor IR6 [Tenebrio molitor] |
| HaxyIR68a.2 | TRINITY_DN6123_c0_g1::TRINITY_DN6123_c0_g1_i1 | 330 | internal | 1 | 417.0 | 5.96e-138 | 58.788 | XP_015839172 | PREDICTED: glutamate receptor ionotropic kainate 5 [Tribolium castaneum] |
| HaxyIR40a.2 | TRINITY_DN8336_c0_g1::TRINITY_DN8336_c0_g1_i1 | 173 | internal | 1 | 259.0 | 4.82e-77 | 65.318 | KYB26635 | hypothetical protein TcasGA2_TC033589 [Tribolium castaneum] |
