## Supplemental tables and figures for "Antennal transcriptome analysis and identification of candidate chemosensory genes of the harlequin ladybird beetle, *Harmonia axyridis* (Pallas) (Coleoptera: Coccinellidae)": Table_S8_OBPs.docx

| Table S8: Best similarity for odorant-binding proteins (OBPs) of *Harmonia axyridis* (Haxy) | | | | | | | | |  |  |  |
| --- | --- | --- | --- | --- | --- | --- | --- | --- | --- | --- | --- |
| Name | ID | length (aa) | ORF status | Signal Peptide | (Total Cysteines) | Conserved Cysteines | Bit score | E-value | % identity | Accession number | Description |
| HaxyOBP1 | TRINITY_DN13152_c0_g1::TRINITY_DN13152_c0_g1_i1 | 140 | 5prime_partial | 0 | 7 | 6 | 222.0 | 1.98e-72 | 73.881 | AJM71483 | odorant-binding protein 9 [Tenebrio molitor] |
| HaxyOBP2 | TRINITY_DN14228_c0_g1::TRINITY_DN14228_c0_g1_i1 | 123 | complete | 1 | 8 | 6 | 52.4 | 3.2e-06 | 31.858 | AIX97058 | odorant-binding protein 12 [Dastarcus helophoroides] |
| HaxyOBP3 | TRINITY_DN16013_c0_g1::TRINITY_DN16013_c0_g1_i1 | 130 | complete | 1 | 8 | 6 | 50.4 | 3e-05 | 50.400 | XP_019876079.1 | PREDICTED: uncharacterized protein LOC109604015 [Aethina tumida] |
| HaxyOBP4 | TRINITY_DN16335_c0_g1::TRINITY_DN16335_c0_g1_i1 | 143 | internal | 0 | 5 | 4 | 75.1 | 1.4e-14 | 37.624 | ALR72510 | odorant binding protein 22 [Colaphellus bowringi] |
| HaxyOBP5 | TRINITY_DN16417_c0_g1::TRINITY_DN16417_c0_g1_i1 | 137 | 5prime_partial | 1 | 6 | 4 | 60.8 | 4.05e-09 | 30.952 | AJM71488 | odorant-binding protein 14 [Tenebrio molitor] |
| HaxyOBP6 | TRINITY_DN16647_c0_g1::TRINITY_DN16647_c0_g1_i1 | 130 | complete | 1 | 6 | 6 | 65.5 | 4.64e-11 | 29.825 | AQY18972 | odorant-binding protein [Galeruca daurica] |
| HaxyOBP7 | TRINITY_DN17055_c0_g1::TRINITY_DN17055_c0_g1_i1 | 135 | 5prime_partial | 1 | 7 | 6 | 64.7 | 1.36e-10 | 32.653 | EFA05742 | odorant binding protein 4 [Tribolium castaneum] |
| HaxyOBP8 | TRINITY_DN17193_c0_g2::TRINITY_DN17193_c0_g2_i1 | 235 | 5prime_partial | 1 | 13 | 6 | 48.5 | 0.001 | 28.000 | XP_017051033.1 | PREDICTED: general odorant-binding protein 56h [Drosophila ficusphila] |
| HaxyOBP9 | TRINITY_DN17357_c0_g1::TRINITY_DN17357_c0_g1_i1 | 138 | 5prime_partial | 1 | 6 | 4 | 52.0 | 8.59e-06 | 24.806 | ARH65461 | odorant binding protein 6 [Anoplophora glabripennis] |
| HaxyOBP10 | TRINITY_DN17574_c0_g1::TRINITY_DN17574_c0_g1_i1 | 143 | complete | 1 | 7 | 6 | 108.0 | 1.5e-27 | 38.889 | AIX97062 | odorant-binding protein 16 [Dastarcus helophoroides] |
| HaxyOBP11 | TRINITY_DN17942_c0_g1::TRINITY_DN17942_c0_g1_i1 | 148 | complete | 1 | 6 | 6 | 211.0 | 4.65e-68 | 65.972 | ALW95359 | odorant-binding protein 2 [Cryptolaemus montrouzieri] |
| HaxyOBP12 | TRINITY_DN17947_c0_g1::TRINITY_DN17947_c0_g1_i1 | 137 | complete | 1 | 4 | 4 | 54.7 | 9.61e-07 | 36.250 | EFA07547 | odorant binding protein C07 [Tribolium castaneum] |
| HaxyOBP13 | TRINITY_DN18004_c0_g1::TRINITY_DN18004_c0_g1_i1 | 152 | complete | 0 | 4 | 4 | 51.6 | 2e-05 | 29.000 | ADO95155.1 | antennal binding protein 7 [Antheraea yamamai] |
| HaxyOBP14 | TRINITY_DN18762_c0_g1::TRINITY_DN18762_c0_g1_i4 | 157 | internal | 0 | 7 | 6 | 252.0 | 4.67e-84 | 85.612 | AVM18960 | odorant binding protein 34 [Holotrichia parallela] |
| HaxyOBP15 | TRINITY_DN18822_c0_g1::TRINITY_DN18822_c0_g1_i3 | 118 | 3prime_partial | 1 | 7 | 6 | 40.0 | 0.21 | 32.000 | AHE13795.1 | odorant binding protein [Lissorhoptrus oryzophilus] |
| HaxyOBP16 | TRINITY_DN19237_c0_g1::TRINITY_DN19237_c0_g1_i1 | 265 | complete | 1 | 14 | 6 | 55.1 | 5.8e-06 | 26.230 | AIX97054 | odorant-binding protein 8 [Dastarcus helophoroides] |
| HaxyOBP17 | TRINITY_DN19337_c0_g1::TRINITY_DN19337_c0_g1_i5 | 101 | 5prime_partial | 0 | 6 | 6 | 172.0 | 1.03e-53 | 91.111 | AVM18963 | odorant binding protein 37, partial [Holotrichia parallela] |
| HaxyOBP18 | TRINITY_DN19353_c0_g1::TRINITY_DN19353_c0_g1_i1 | 174 | complete | 1 | 9 | 6 | 245.0 | 1.82e-80 | 68.571 | XP_008194712 | PREDICTED: general odorant-binding protein 70 isoform X2 [Tribolium castaneum]<>hypothetical protein TcasGA2_TC016310 [Tribolium castaneum] |
| HaxyOBP19 | TRINITY_DN19563_c0_g1::TRINITY_DN19563_c0_g1_i4 | 140 | complete | 1 | 5 | 4 | 49.0 | 1e-04 | 26.000 | EFA07429.1 | odorant binding protein C09 [Tribolium castaneum] |
| HaxyOBP20 | TRINITY_DN19737_c0_g2::TRINITY_DN19737_c0_g2_i3 | 145 | complete | 1 | 6 | 6 | 240.0 | 1.01e-79 | 81.884 | AVM18959 | odorant binding protein 33, partial [Holotrichia parallela] |
| HaxyOBP21 | TRINITY_DN19794_c0_g3::TRINITY_DN19794_c0_g3_i4 | 137 | 3prime_partial | 1 | 7 | 6 | 190.0 | 8.79e-60 | 63.910 | ALW95359 | odorant-binding protein 2 [Cryptolaemus montrouzieri] |
| HaxyOBP22 | TRINITY_DN19794_c0_g4::TRINITY_DN19794_c0_g4_i1 | 148 | complete | 1 | 7 | 6 | 246.0 | 1.21e-81 | 79.730 | ALW95359 | odorant-binding protein 2 [Cryptolaemus montrouzieri] |
| HaxyOBP23 | TRINITY_DN20561_c0_g1::TRINITY_DN20561_c0_g1_i1 | 127 | complete | 1 | 5 | 4 | 122.0 | 1.48e-33 | 48.092 | XP_975685 | PREDICTED: B1 protein [Tribolium castaneum]<>odorant binding protein (subfamily minus-C) C04 [Tribolium castaneum] |
| HaxyOBP24 | TRINITY_DN20835_c2_g3::TRINITY_DN20835_c2_g3_i5 | 103 | 5prime_partial | 0 | 6 | 6 | 43.9 | 0.006 | 34.000 | XP_015836450.1 | PREDICTED: general odorant-binding protein 57c [Tribolium castaneum] |
| HaxyOBP25 | TRINITY_DN21686_c2_g1::TRINITY_DN21686_c2_g1_i1 | 126 | 5prime_partial | 1 | 6 | 4 | 47.8 | 4e-04 | 31.000 | AIX97054.1 | odorant-binding protein 8 [Dastarcus helophoroides] |
| HaxyOBP26 | TRINITY_DN22387_c0_g2::TRINITY_DN22387_c0_g2_i1 | 184 | 3prime_partial | 1 | 9 | >6 | 188.0 | 7.37e-57 | 51.872 | XP_015836161 | PREDICTED: uncharacterized protein LOC660933 [Tribolium castaneum]<>hypothetical protein TcasGA2_TC013160 [Tribolium castaneum] |
| HaxyOBP27 | TRINITY_DN22446_c0_g1::TRINITY_DN22446_c0_g1_i1 | 144 | 5prime_partial | 1 | 4 | 4 | 153.0 | 9.78e-46 | 75.258 | AVM18964 | odorant binding protein 38, partial [Holotrichia parallela] |
| HaxyOBP28 | TRINITY_DN22673_c0_g2::TRINITY_DN22673_c0_g2_i1 | 152 | 5prime_partial | 0 | 4 | 4 | 52.4 | 9.73e-06 | 31.538 | APG79376 | pheromone binding protein 15 [Cyrtotrachelus buqueti] |
| HaxyOBP29 | TRINITY_DN22984_c0_g1::TRINITY_DN22984_c0_g1_i1 | 151 | complete | 1 | 7 | 6 | 62.0 | 1.76e-09 | 28.058 | AIX97054 | odorant-binding protein 8 [Dastarcus helophoroides] |
| HaxyOBP30 | TRINITY_DN23033_c0_g1::TRINITY_DN23033_c0_g1_i1 | 220 | complete | 1 | 8 | 6 | 126.0 | 1.96e-32 | 32.489 | AGI05159 | odorant-binding protein 21 [Dendroctonus ponderosae]<>hypothetical protein D910_01855 [Dendroctonus ponderosae] |
| HaxyOBP31 | TRINITY_DN23312_c0_g1::TRINITY_DN23312_c0_g1_i1 | 135 | complete | 1 | 6 | 6 | 52.8 | 4.54e-06 | 24.390 | AIX97054 | odorant-binding protein 8 [Dastarcus helophoroides] |
