## Supplemental tables and figures for "Antennal transcriptome analysis and identification of candidate chemosensory genes of the harlequin ladybird beetle, *Harmonia axyridis* (Pallas) (Coleoptera: Coccinellidae)": Table_S9_CSPs.docx

| Table S9: Best similarity for chemosensory proteins (CSPs) of *Harmonia axyridis* (Haxy) | | | | | | | | |  |  |  |
| --- | --- | --- | --- | --- | --- | --- | --- | --- | --- | --- | --- |
| Name | ID | length (aa) | ORF status | Signal Peptide | (Total Cysteines) | Conserved Cysteines | Bit score | E-value | % identity | Accession number | Description |
| HaxyCSP1 | TRINITY_DN14407_c0_g1::TRINITY_DN14407_c0_g1_i1 | 134 | 5prime_partial | 1 | 6 | 4 | 140.0 | 3.31e-40 | 47 | NP_001039278 | chemosensory protein 10 precursor [Tribolium castaneum]<>chemosensory protein 10 [Tribolium castaneum]<>chemosensory protein 7 [Tribolium castaneum] |
| HaxyCSP2 | TRINITY_DN16195_c0_g1::TRINITY_DN16195_c0_g1_i1 | 108 | complete | 1 | 4 | 4 | 161.0 | 2.44e-49 | 73 | AJO62216 | chemosensory protein CSP10 [Tenebrio molitor] |
| HaxyCSP3 | TRINITY_DN16996_c0_g1::TRINITY_DN16996_c0_g1_i2 | 127 | complete | 1 | 5 | 4 | 157.0 | 2.19e-47 | 71 | NP_001039280 | chemosensory protein 12 precursor [Tribolium castaneum]<>chemosensory protein 12 [Tribolium castaneum]<>chemosensory protein 9 [Tribolium castaneum] |
| HaxyCSP4 | TRINITY_DN17208_c0_g1::TRINITY_DN17208_c0_g1_i1 | 132 | complete | 1 | 5 | 4 | 175.0 | 2.71e-54 | 63 | NP_001039289 | chemosensory protein 7 precursor [Tribolium castaneum]<>PREDICTED: chemosensory protein 7 isoform X1 [Tribolium castaneum]<>chemosensory protein 7 [Tribolium castaneum]<>chemosensory protein 12 [Tribolium castaneum] |
| HaxyCSP5 | TRINITY_DN18656_c0_g3::TRINITY_DN18656_c0_g3_i1 | 140 | 5prime_partial | 1 | 6 | 4 | 157.0 | 3.99e-47 | 53 | NP_001039278 | chemosensory protein 10 precursor [Tribolium castaneum]<>chemosensory protein 10 [Tribolium castaneum]<>chemosensory protein 7 [Tribolium castaneum] |
| HaxyCSP6 | TRINITY_DN18685_c2_g1::TRINITY_DN18685_c2_g1_i1 | 129 | complete | 1 | 4 | 4 | 133.0 | 1.25e-37 | 54 | XP_011202324 | PREDICTED: ejaculatory bulb-specific protein 3 [Bactrocera dorsalis]<>chemosensory protein 2 [Bactrocera dorsalis] |
| HaxyCSP7 | TRINITY_DN19896_c0_g1::TRINITY_DN19896_c0_g1_i1 | 101 | complete | 1 | 5 | 4 | 121.0 | 7.99e-34 | 62 | AIX97044 | chemosensory protein 4 [Monochamus alternatus] |
| HaxyCSP8 | TRINITY_DN20287_c0_g2::TRINITY_DN20287_c0_g2_i3 | 140 | complete | 1 | 4 | 4 | 169.0 | 1.06e-51 | 60 | AIX97040 | chemosensory protein 8 [Monochamus alternatus] |
| HaxyCSP9 | TRINITY_DN20882_c0_g1::TRINITY_DN20882_c0_g1_i1 | 153 | complete | 0 | 6 | 4 | 154.0 | 1.74e-45 | 55 | AKK25146 | chemosensory protein 1 [Dendroctonus ponderosae] |
| HaxyCSP10 | TRINITY_DN22918_c0_g2::TRINITY_DN22918_c0_g2_i5 | 124 | complete | 1 | 5 | 4 | 137.0 | 2.43e-39 | 54 | CAJ01490 | hypothetical protein [Biphyllus lunatus] |
| HaxyCSP11 | TRINITY_DN7721_c0_g1::TRINITY_DN7721_c0_g1_i1 | 103 | internal | 0 | 4 | 4 | 202.0 | 1.95e-65 | 97 | AGE97642 | chemosensory protein 2 [Aphis gossypii] |
| HaxyCSP12 | TRINITY_DN8934_c0_g1::TRINITY_DN8934_c0_g1_i1 | 145 | 5prime_partial | 0 | 5 | 4 | 160.0 | 4.16e-48 | 57 | NP_001039278 | chemosensory protein 10 precursor [Tribolium castaneum]<>chemosensory protein 10 [Tribolium castaneum]<>chemosensory protein 7 [Tribolium castaneum] |
