## Supplemental tables and figures for "Antennal transcriptome analysis and identification of candidate chemosensory genes of the harlequin ladybird beetle, *Harmonia axyridis* (Pallas) (Coleoptera: Coccinellidae)": Table_S10_SNMPs.docx

| Table S10: Best similarity for sensory neuron membrane proteins (SNMPs) of *Harmonia axyridis* (Haxy) | | | | | | | | | |
| --- | --- | --- | --- | --- | --- | --- | --- | --- | --- |
| Name | ID | length (aa) | ORF status | TMHMM | Bit score | E-value | % identity | Accession number | Description |
| HaxySNMP1.1 | TRINITY_DN24414_c1_g1::TRINITY_DN24414_c1_g1_i2 | 525 | 3prime_partial | 2 | 688 | 0 | 62.500 | XP_001816436 | PREDICTED: sensory neuron membrane protein 1 [Tribolium castaneum] |
| HaxySNMP1.2 | TRINITY_DN24697_c2_g1::TRINITY_DN24697_c2_g1_i1 | 554 | 5prime_partial | 2 | 533 | 0 | 51.329 | ALR72543 | sensory neuron membrane protein SNMP1b [Colaphellus bowringi] |
| HaxySNMP2.1 | TRINITY_DN5381_c0_g1::TRINITY_DN5381_c0_g1_i1 | 205 | 5prime_partial | 1 | 270 | 1.54e-85 | 63.636 | XP_008198962 | PREDICTED: sensory neuron membrane protein 2 [Tribolium castaneum] |
| HaxySNMP2.2 | TRINITY_DN5381_c0_g2::TRINITY_DN5381_c0_g2_i1 | 207 | 3prime_partial | 1 | 216 | 9.94e-65 | 49.029 | XP_008198962 | PREDICTED: sensory neuron membrane protein 2 [Tribolium castaneum] |
