## Supplementary figures and images for "Antennal transcriptome analysis and identification of candidate chemosensory genes of the harlequin ladybird beetle, *Harmonia axyridis* (Pallas) (Coleoptera: Coccinellidae)"

### Figure_S1.png

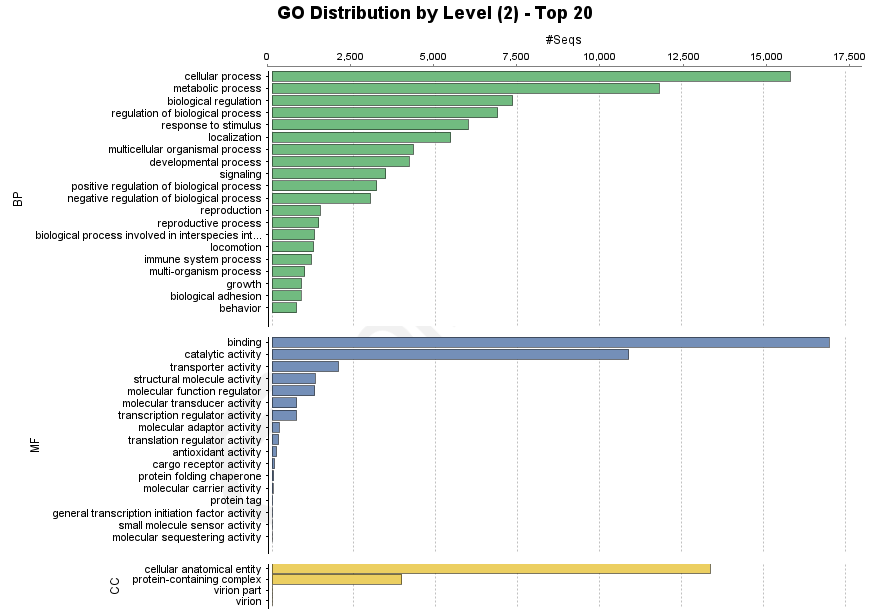

### Figure_S2.pdf

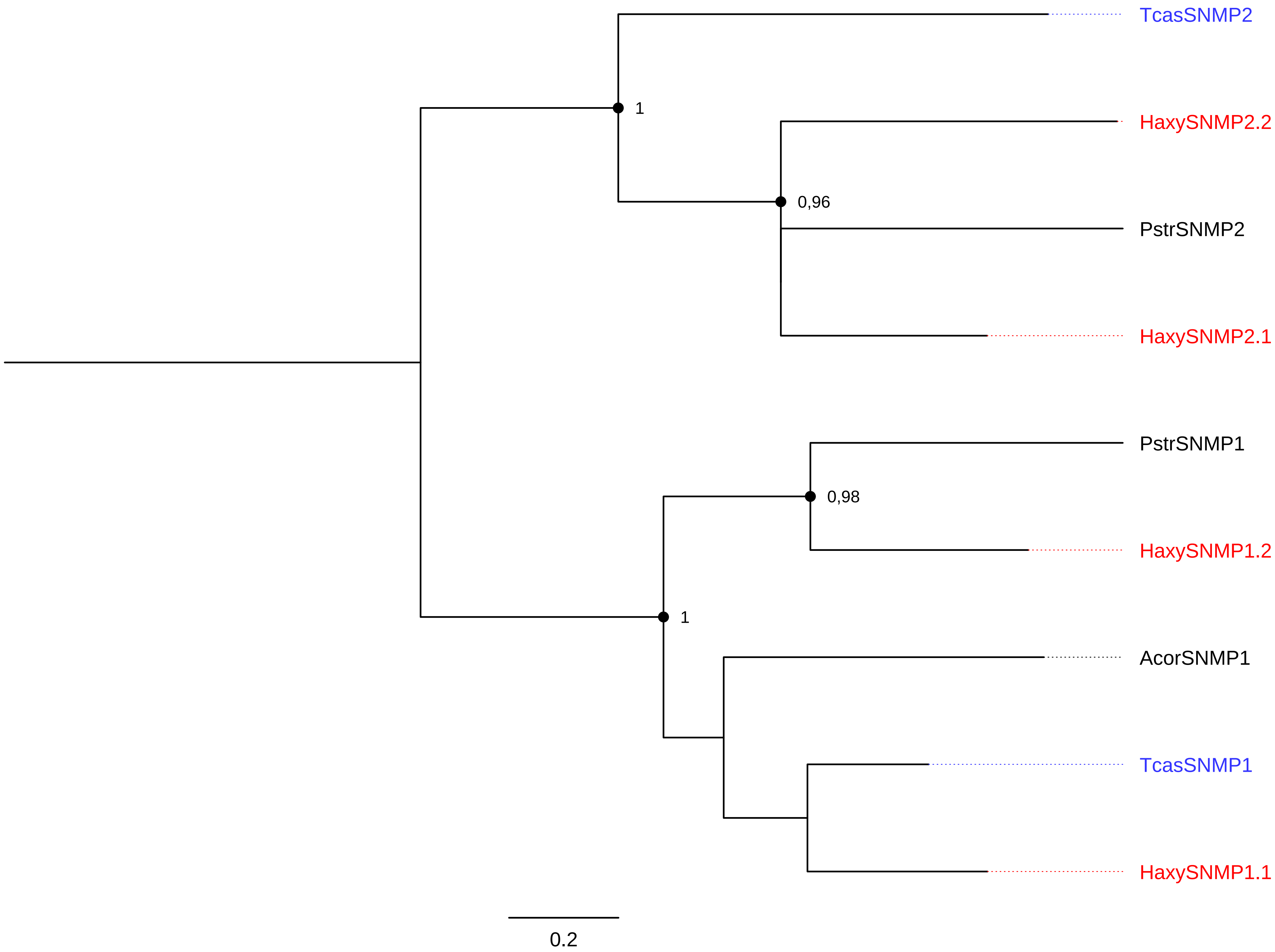
